# V3Cell: A Vision-Guided Virtual 3D Cell Framework for Phenotypic Modeling and Perturbation Prediction

**DOI:** 10.64898/2026.06.23.734130

**Authors:** Lu You, Xun Deng, Chenke Xu, Xiaobo Zhu, Zhigang Zhang, Pengyu Chen, Xiwen Yang, Zhengzheng Yan, Jiahua Rao, Huili Hu, Jianying Hu, Pengwei Hu

## Abstract

Predicting how organoids respond to chemical perturbations is central to disease modeling and drug discovery. Existing virtual cell models operate at the single-cell level, producing static endpoint predictions from destructive assays. This leaves a critical gap at the organoid scale, where biological identity is defined by tissue-level architecture and continuous developmental dynamics rather than single-cell features. Here we introduce V3Cell, a vision-guided framework that constructs in silico surrogates of organoids directly from non-invasive bright-field microscopy. A foreground-aware model constructs static virtual 3D cells across colon, stomach, and lung organoid lineages. These virtual 3D cells closely match real samples across distributional metrics, micro-texture, and lineage-specific morphometrics, with small effect sizes for most descriptors. A temporal module further predicts developmental fate from as few as six early-frame observations and models fate-conditioned spatiotemporal trajectories that closely recapitulate real perturbation responses. V3Cell requires no omics profiling or fluorescent labeling, establishing a non-invasive brightfield-based paradigm for organoid-scale perturbation prediction. Our code and data are publicly available at https://github.com/Laineyoulu/V3Cell.

## 1 Introduction

Three-dimensional cell constructs self-organize from living cells and exhibit emergent structural and functional properties that are absent in conventional two-dimensional cultures. Organoids recapitulate this three-dimensional cellular organization in vitro, faithfully reproducing the architecture and physiology of their parent organs [1, 2]. A central challenge in disease modeling and drug development is to understand and predict how organoids respond to external perturbations, such as drug treatments [3, 4]. The ability to predict these organoid state changes in silico would enable large-scale pre-screening of candidate compounds before committing to wet-lab validation, reducing the time and cost of experimental screening [5, 6]. The concept of the virtual cell, which leverages computational models to simulate cellular behavior, has emerged as a promising paradigm to address this challenge [7–11].

Driven by advances in deep learning and single-cell sequencing, a growing family of virtual cell models has been developed for perturbation prediction [12, 13]. The dominant paradigm operates in molecular feature space, where models such as scGen [14], CPA [15], GEARS [16], and CellOT [17] learn to map perturbation contexts directly to post-perturbation gene expression profiles. Meanwhile, foundation models including scGPT [18], scFoundation [19], and Geneformer [20] leverage large-scale pretraining to adapt to perturbation tasks. Collectively, these molecular-space methods operate at the single-cell level and produce static, endpoint predictions of transcriptomic states. More recently, image-based generative approaches such as CellFlux [21], MorphGen [22], MorphoDiff [23], and PhenDiff [24] have begun to model perturbation-induced morphological changes at the single-cell level[25, 26], typically taking multichannel fluorescent Cell Painting images as their input and output modalities. These image-based methods share the same single-cell scope as their molecular-space counterparts and typically depend on multichannel fluorescent staining for data acquisition.

Despite this progress, fundamental gaps remain when the modeling target is extended from isolated single cells to complex 3D multicellular systems [27, 28]. From a modeling perspective, both molecular-space and existing image-based approaches operate at the single-cell level and produce predictions at static perturbation end-points. Although some methods support smooth interpolation between cell states [17, 29], few, if any, predict temporally ordered frame sequences at the organoid scale, conditioned on brightfield observations alone and without molecular priors. A separate body of work has developed non-invasive tools for brightfield-based organoid segmentation and longitudinal tracking [30–32], but these tools aim to quantify observed trajectories rather than to forecast future ones. This distinction is critical at the organoid scale, because organoid phenotypic identity arises from the collective self-organization of thousands of cells into spatially structured architectures [33, 34], and its morphology is defined by tissue-level architecture, internal texture, and three-dimensional context. Existing single-cell models are designed to capture single-cell-level features by construction [11], whereas capturing these tissue-level emergent features requires the model to ingest and model organoid-scale observations directly [35, 36]. This requirement points naturally toward image-based modeling at the whole-organoid level [37, 38].

From a data perspective, current approaches depend on modalities that are fundamentally incompatible with longitudinal observation. Transcriptomic profiling requires cell dissociation and lysis [39–42], while fluorescence-based cell painting demands fixation and staining [43–45]. Both are destructive, precluding repeated measurement of the same specimen and making it difficult to track the continuous developmental trajectories that define organoid biology [46, 47]. While live fluorescent reporters partially mitigate this, they require transgenic labeling and are restricted to a narrow set of markers, limiting the biological scope that can be observed longitudinally. For organoids, this constraint is further amplified. Comprehensive molecular or fluorescence characterization demands costly protocols and labor-intensive preparation. More critically, molecular profiling often destroys the intact 3D architecture of the specimen, whereas endpoint fluorescence assays typically preclude continued longitudinal observation of the same living organoid.

Constructing a virtual surrogate for organoids therefore demands a fundamentally different approach. Such a framework must instantiate an observable entity with multi-scale spatial structure and continuous temporal dynamics, and it must derive this entity from a modality that is both non-invasive and longitudinally compatible. Bright-field microscopy provides a practical compromise that meets these requirements. As a stain-free imaging modality, it captures morphological architecture, internal texture, and spatial organization without fixation, staining, or perturbation of the specimen [48–51]. Because the specimen remains intact and viable, brightfield imaging supports continuous time-lapse recording of individual organoids over extended culture periods [52, 53]. This produces specimen-anchored developmental trajectories that are inaccessible to any destructive assay. Moreover, visual phenotyping is not merely a convenient proxy. The molecular and cellular states that define organoid identity ultimately manifest as observable morphological features, establishing brightfield imaging as an empirically supported readout of the underlying biological state [54–57].

Building on these principles, we introduce V3Cell, a vision-guided modeling framework for the construction of virtual 3D cells, which are in silico surrogates of organoids constructed directly from 2D brightfield microscopy (Figure 1). V3Cell adopts a two-stage architecture. The first stage learns latent representations of organoid visual phenotypes at the region-of-interest (ROI) level, where each ROI is a single organoid cropped from the imaging field rather than the entire field itself. This stage also enables controllable construction across diverse organoid lineages and morphological categories. Because organoid imaging fields are dominated by background artifacts such as Matrigel, debris, and out-of-focus neighboring organoids[30, 58], a foreground-aware learning strategy directs the model to prioritize organoid-centric morphological features while suppressing background confounds. The second stage extends the framework to the temporal domain. Conditioned on sparse early-frame observations, V3Cell first predicts the organoid developmental fate and then predicts fate-conditioned spatiotemporal trajectories of its morphological evolution and treatment response, where the spatial dimension refers to 2D image-plane morphology. By operating entirely within the observational modality of standard brightfield microscopy, V3Cell requires neither omics profiling nor destructive endpoints, establishing a non-invasive brightfield-based paradigm for in silico virtual 3D cell construction and perturbation prediction. We highlight three main contributions of our study:

**Fig. 1.**
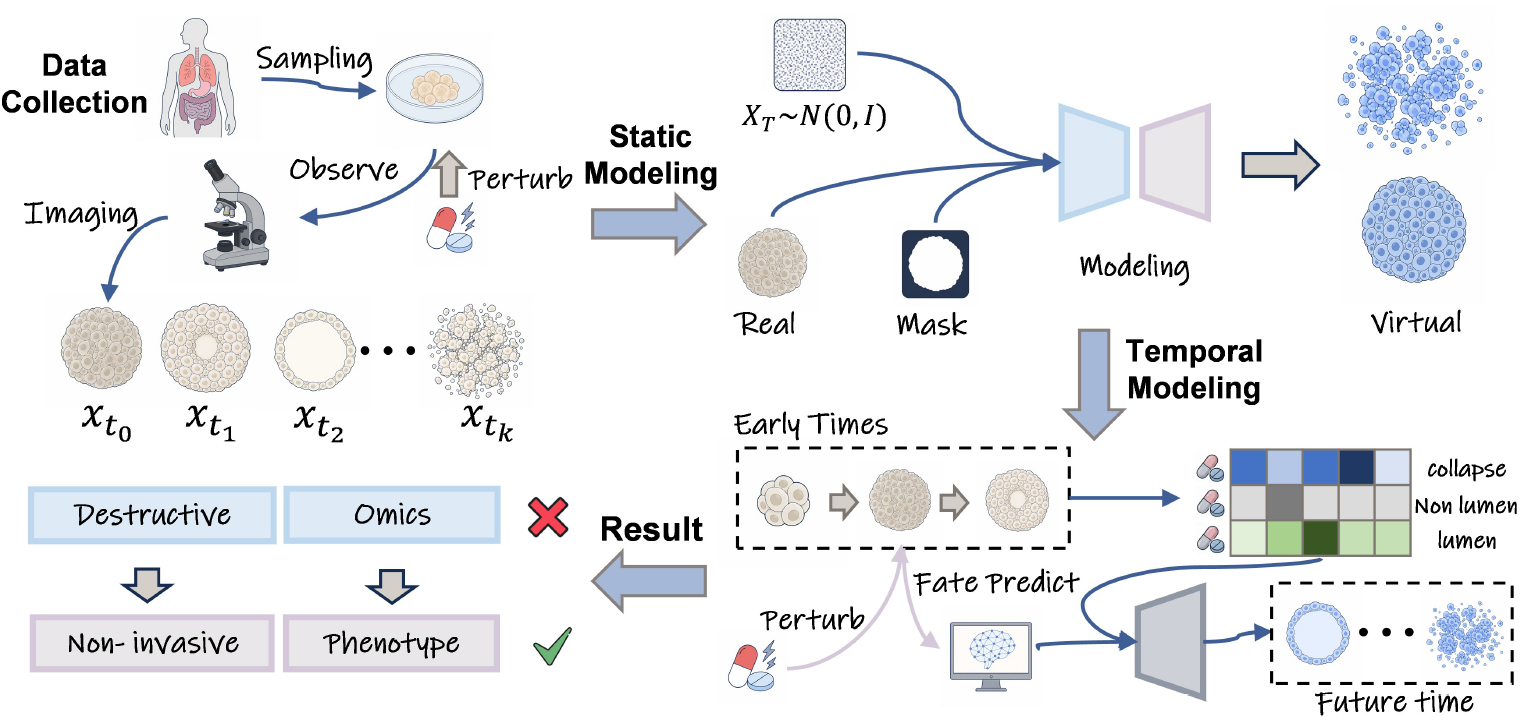
Overview of the V3Cell framework for non-invasive virtual 3D cell modeling and perturbation prediction. **Data collection**. Brightfield time-lapse sequences of organoids are acquired under baseline and chemically perturbed conditions, yielding longitudinal observa-tions 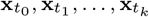 . This label-free imaging modality preserves specimen viability and enables repeated phenotypic observation without destructive omics profiling. **Stage 1: Static modeling**. A foreground-aware conditional model constructs high-fidelity static virtual 3D cells from sampling noise and phenotype-conditioning information. During training, foreground masks derived from real organoid images impose organoid-centered constraints. At inference, masks are not provided. **Stage 2: Temporal modeling and fate prediction**. Conditioned on sparse early-frame observations, a temporal modeling module first predicts organoid developmental fate and then autoregressively forecasts fate-conditioned future morphological trajectories, producing dynamic virtual 3D cells that capture spatiotemporal evolution under perturbation.

- **Establishing the Virtual 3D Cell Paradigm**. We introduce the concept of virtual 3D cells as in silico surrogates of 3D multicellular systems that are constructed as observable entities with image-plane spatial structure and temporal dynamics. This complements existing models that operate at the single-cell level with static, endpoint predictions[7, 11] by extending the paradigm to the organoid scale. V3Cell realizes this paradigm by learning latent representations of organoid states directly from brightfield images, providing a non-destructive, scalable alternative to both omics-based and fluorescence-based modeling.
- **High-Fidelity Construction with ROI-Level Conditioning**. We demonstrate that V3Cell constructs high-quality static virtual 3D cells across diverse organoid categories and culture contexts. Through a foreground-aware learning strategy, the model supports class-conditioned ROI-level construction of visual phenotypes while suppressing Matrigel, debris, and out-of-focus background. These static representations serve as a foundation for longitudinal modeling and morphological analysis.
- **Longitudinal Fate Prediction and Perturbation Modeling of Virtual 3D Cells**. We show that V3Cell supports predictive modeling of organoid fate under external perturbations using as few as *K* = 6 early-frame observations. By first predicting the developmental fate from these sparse observations and then conditioning on the predicted fate, V3Cell predicts ROI-level spatiotemporal trajectories of future developmental outcomes and treatment responses, enabling early phenotypic response forecasting from sparse imaging data alone.

## 2 Results

### 2.1 V3Cell constructs a high-quality static virtual 3D cell resource

To assess whether brightfield microscopy alone provides sufficient signal for phenotype-faithful virtual 3D cell construction, we applied V3Cell across three organoid lineages encompassing a broad range of morphological subtypes. The constructed virtual 3D cells faithfully recapitulated lineage-specific hallmark morphologies across all systems, spanning colon organoids stratified by budding status and optical transparency, stomach organoids with apoptotic, hollow, or solid architectures, and lung organoids with lumen or non-lumen organization (Fig. 2). V3Cell modeled both canonical phenotypic archetypes and visually intermediate states within the annotated categories. Reproducible construction across phenotypic categories indicates that the learned representation organizes major visual phenotypes consistently. Together, these results indicate that non-invasive brightfield imaging encodes sufficient morphological information to support high-throughput virtual 3D cell construction across diverse organoid systems, providing the static virtual-cell basis for the distributional, morphometric, and phenotyping analyses below.

**Fig. 2.**
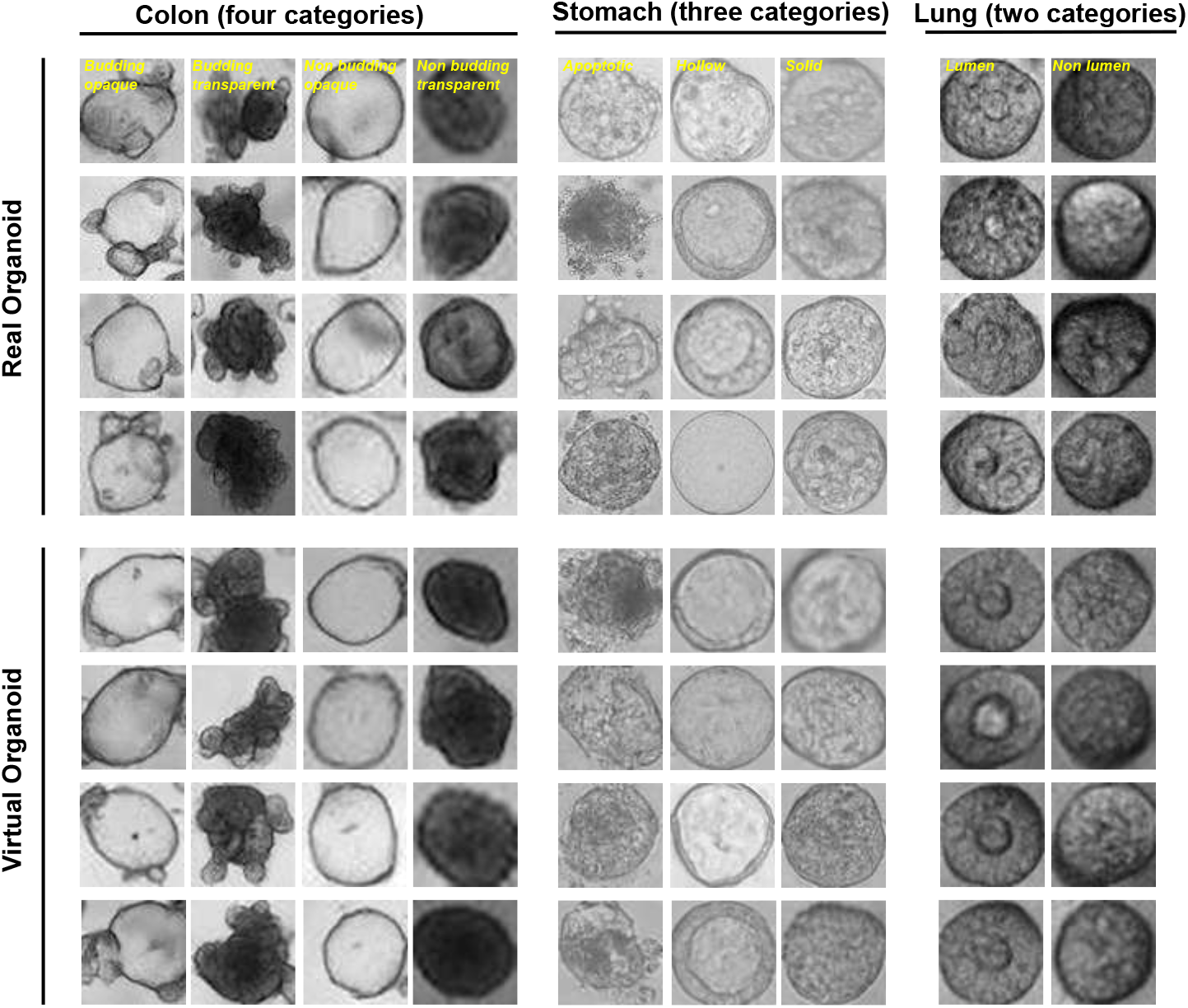
Comparison of real organoids and V3Cell-constructed virtual 3D cells across diverse phenotypes. Representative brightfield images of real organoids are shown in the upper block, and category-matched virtual 3D cells constructed by V3Cell are shown in the lower block. Samples are organized by organoid lineage and phenotype class, including four colon categories, three stomach categories, and two lung categories as labeled in the figure. Virtual examples were modeled using a fixed random seed for display consistency.

### 2.2 Multi-dimensional evaluation of virtual 3D cell quality

To validate V3Cell as a reliable in silico model, we assessed whether the constructed virtual 3D cells statistically recapitulate the population-level distributions of real samples. We benchmarked V3Cell against representative generative baselines, including conditional GANs (cGAN[59], F2DGAN[60]), a conditional VAE (cVAE[61]), and diffusion models (LDM[62], DiT[63], Medfusion[64]). To quantify distributional fidelity, we employed the Inception Score (IS) and Kernel Inception Distance (KID) (formulations in Methods and Table S1). Across all organoid systems, V3Cell achieved the highest IS and lowest KID (Fig. 3A), indicating strong alignment with the real phenotypic manifold. On the colon dataset, V3Cell recorded a low KID of 1.9 × 10^−5^ ± 1.0 × 10^−6^, indicating a small discrepancy between real and virtual feature distributions. V3Cell attained the lowest KID across all three organoid lineages, reducing KID by up to 2.7-fold relative to the best-performing baseline, and the highest IS in all systems, indicating consistent advantages in both distributional alignment and phenotypic diversity across distinct biological contexts.

**Fig. 3.**
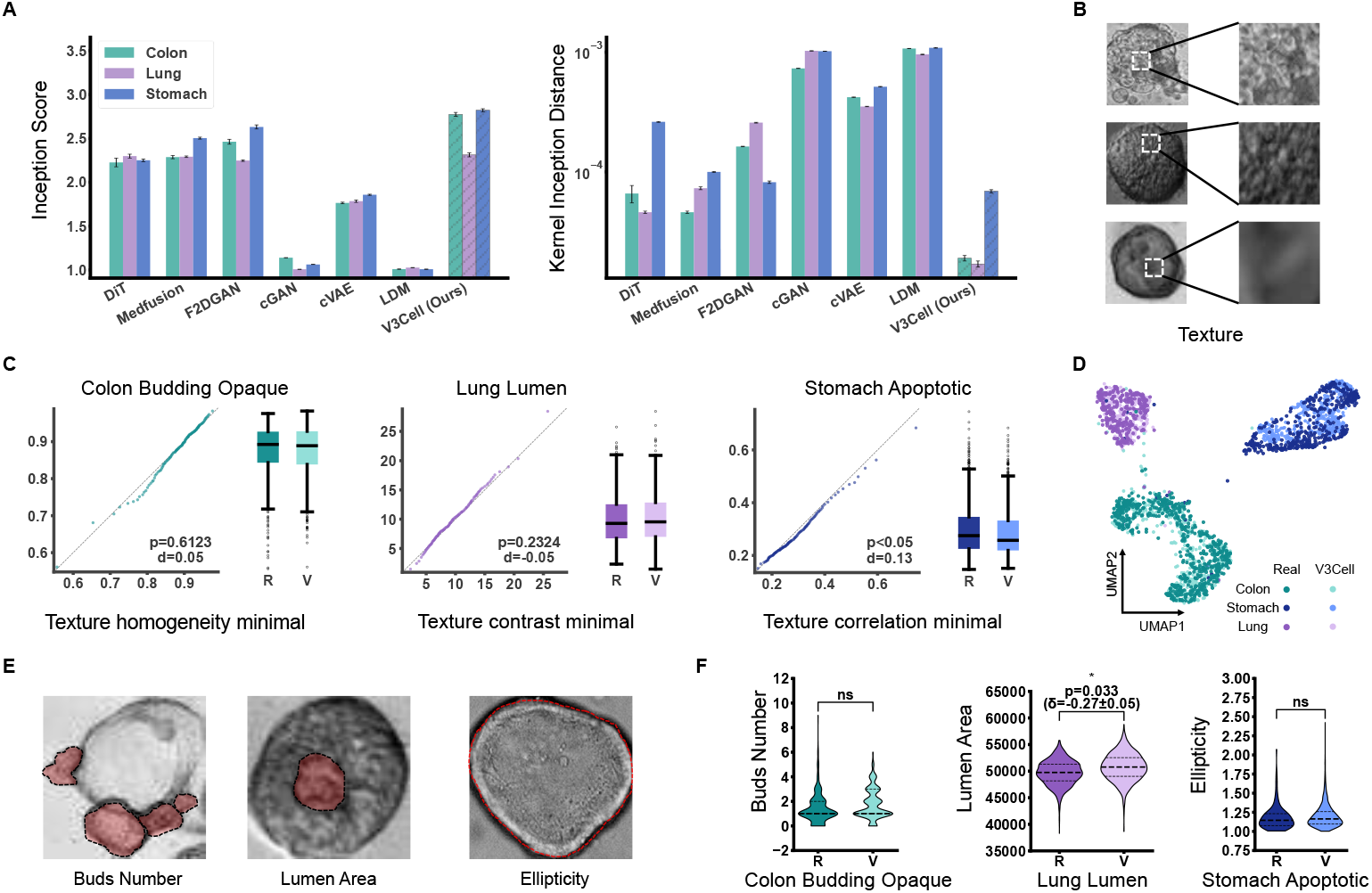
Multi-dimensional evaluation of virtual 3D cell quality and phenotypic fidelity. **(A)** Quantitative benchmarking of V3Cell against generative baselines including DiT, Medfusion, F2DGAN, cGAN, cVAE, and LDM. Bar charts show Inception Score and Kernel Inception Distance across colon, lung, and stomach datasets. **(B)** Representative brightfield micrographs of real organoids with magnified regions highlighting local micro-textural patterns. **(C)** Micro-textural fidelity evaluated using representative Haralick descriptors for Colon Budding Opaque, Lung Lumen, and Stomach Apoptotic organoids. Q-Q plots show quantile alignment between real organoids and virtual 3D cells, and box plots show the corresponding marginal distributions. **(D)** UMAP projection of high-dimensional organoid features from real and V3Cell-constructed populations across colon, lung, and stomach lineages. **(E)** Visual annotations of lineage-specific morphometric hallmarks, including bud number in colon, lumen area in lung, and ellipticity in stomach organoids. **(F)** Violin plots comparing morphometric distributions between real organoids and virtual 3D cells for Colon Budding Opaque, Lung Lumen, and Stomach Apoptotic organoids. Statistical significance was assessed by Wilcoxon rank-sum test, with effect sizes reported as Cohen’s *d±* s.e.

While distributional metrics capture population-level fidelity, they do not directly assess whether virtual 3D cells preserve the fine-grained micro-textural properties that distinguish different organoid subtypes. As shown in Fig. 3B, distinct organoid sub-types exhibit characteristic micro-textural patterns. We therefore quantified Haralick texture descriptors [65], including contrast, correlation, homogeneity, entropy, and Information Measure of Correlation 1 (IMC1), across representative subtypes from each lineage (Fig. 3C), with the complete set of evaluations provided in Supplementary Fig. S1. Given the large sample sizes employed, we focused on effect size rather than statistical significance as the primary measure of practical difference. Cohen’s *d* [66] values remained below 0.2 across most metrics and organoid subtypes, indicative of a small effect size between real and virtual cell distributions. Quantile-quantile plots showed tight alignment with the identity line across evaluated subtypes, supporting close agreement between real and virtual organoids in micro-textural statistics.

To integrate these multi-scale descriptors into a unified phenotypic landscape, we projected high-dimensional feature vectors from our organoid-specific encoder using Uniform Manifold Approximation and Projection (UMAP) [67]. Virtual 3D cell populations were embedded within and overlapped extensively with the manifold of their real counterparts across all lineages (Fig. 3D), supporting broad coverage of the real phenotypic manifold and showing no evidence of gross mode collapse.

The biological fidelity of this virtual space is further evidenced by the precise recapitulation of lineage-specific developmental hallmarks (Fig. 3E–F, Supplementary Fig. S2). In colon budding opaque organoids, V3Cell reproduced the distribution of bud number, a critical emergent feature of colon crypt formation, with no significant difference from real samples. In lung lumen organoids, the modeled lumen area closely aligned with real observations, showing only a small residual difference that was negligible in effect size (*p* = 0.033, *d* = −0.27 ±0.05). For stomach apoptotic organoids, ellipticity distributions showed no significant difference between real and virtual cells, indicating that our foreground-aware training strategy effectively prioritizes organoid-centric features. Collectively, these results establish that virtual 3D cells are not merely visually plausible but are closely aligned with real organoids at the ROI level, with small effect sizes across descriptors, providing a high-fidelity foundation for the longitudinal analyses that follow.

### 2.3 Virtual 3D cells empower reliable phenotyping

Beyond statistical realism, we asked whether virtual 3D cells provide functional utility for downstream phenotyping tasks under the sparse sampling constraints typical of longitudinal organoid studies. All evaluations were conducted exclusively from brightfield images, without omics profiling or fluorescence labeling.

We first assessed whether virtual 3D cells alone encode sufficient biological information for phenotypic discrimination. A classifier trained exclusively on V3Cell-constructed virtual 3D cells, without any physical samples, achieved macro-*F*_1_ scores of 0.86 in colon, 0.78 in stomach, and 0.82 in lung organoids, indicating that virtual cells preserve lineage-specific morphological information sufficient for phenotypic discrimination.

We then simulated sparse physical sampling constraints. Relying solely on 25 physical samples per category caused macro-*F*_1_ scores to degrade to 65.8% in colon, 59.4% in stomach, and 31.7% in lung organoids (Fig. 4A). Augmenting these limited sets with V3Cell-constructed virtual 3D cells restored performance to 87.8%, 83.4%, and 76.4%, respectively, yielding improvements of 22.0, 24.1, and 44.7 percentage points (Fig. 4B). This augmentation effect was consistent across a broader range of low-resource regimes spanning 10 to 50 samples per class, with the most pronounced recovery occurring in the lung lineage.

**Fig. 4.**
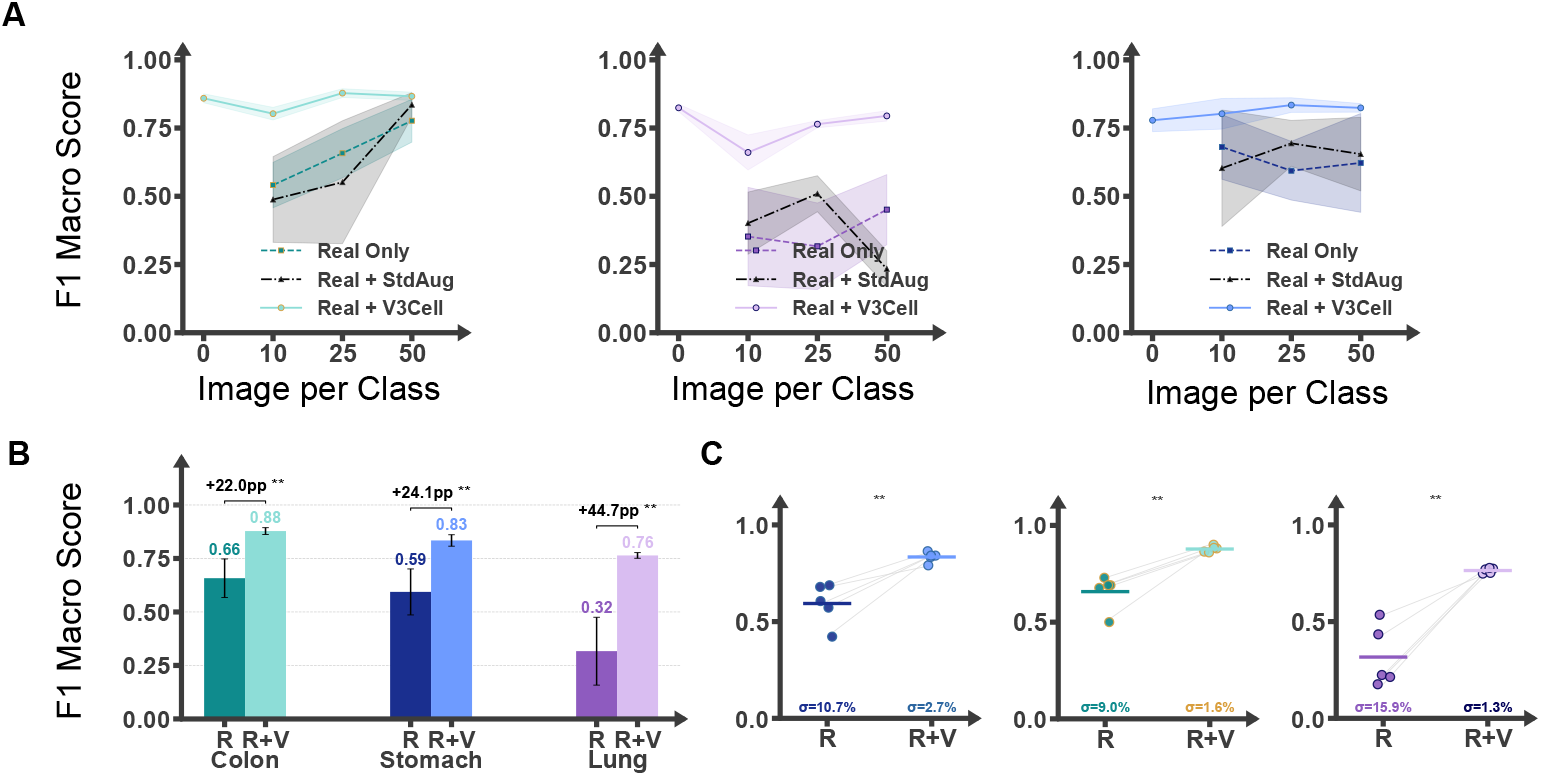
Virtual 3D cells empower reliable phenotyping in low-resource regimes. **(A)** Phenotypic classification performance (macro-*F*_1_) across varying quantities of physical training samples (10, 25, and 50 per class) for colon, stomach, and lung organoids, comparing real-only, standard augmentation, and V3Cell augmentation conditions. Shaded regions indicate *±*1 standard deviation across *n* = 5 independent random seeds. **(B)** Quantitative comparison of augmentation effect at 25 physical samples per class across all three lineages. Error bars denote standard deviations. **(C)** Phenotyping reproducibility at 25 samples per class. Lines connect paired experimental runs sharing identical physical training splits, demonstrating that V3Cell augmentation consistently reduces inter-run variance and stabilizes classification performance.

Beyond classification accuracy, virtual augmentation substantially reduced interrun variability. At 25 samples per class, the real-only lung classifier exhibited a standard deviation of 15.9% in macro-*F*_1_ score. Incorporating virtual 3D cells reduced this variance to 1.3% (Fig. 4C), indicating that virtual augmentation can stabilize phenotypic decision boundaries under sparse physical sampling conditions.

### 2.4 V3Cell predicts organoid fate trajectories and perturbation responses from sparse early frames

To evaluate whether V3Cell can recapitulate fate-specific morphological trajectories and perturbation responses from sparse conditioning frames, we designed a systematic quantitative assessment spanning representation learning, early-frame classification, and trajectory fidelity. For this longitudinal analysis we employed an independent brightfield time-lapse dataset of hESC-derived amniotic-sac organoids (rosette-like 3D structures) undergoing apical cavity formation under chemical perturbation, which is biologically and experimentally distinct from the static colon, stomach, and lung organoid datasets analyzed in Sections 2.1–2.3 (dataset details in Methods). Within this assay we focused on three distinct developmental fate classes: cavity formation, cavity collapse, and non-cavity outcome. Each fate class represents a characteristic developmental endpoint whose probability is modulated by external chemical perturbation. Importantly, neither fate prediction nor trajectory modeling requires knowledge of the applied perturbation at inference time. Both are driven entirely by morphological features extracted from early brightfield observations, and the predicted fate label then serves as the sole categorical conditioning signal supplied to the frozen image generator that renders each predicted frame (Fig. 5A). Notably, across this OPFR family, developmental toxicity could neither be separated by molecular structure nor predicted from it at screening-relevant accuracy (Supplementary Figs. S4 and S5), motivating the phenotype-based modeling used in V3Cell.

**Fig. 5.**
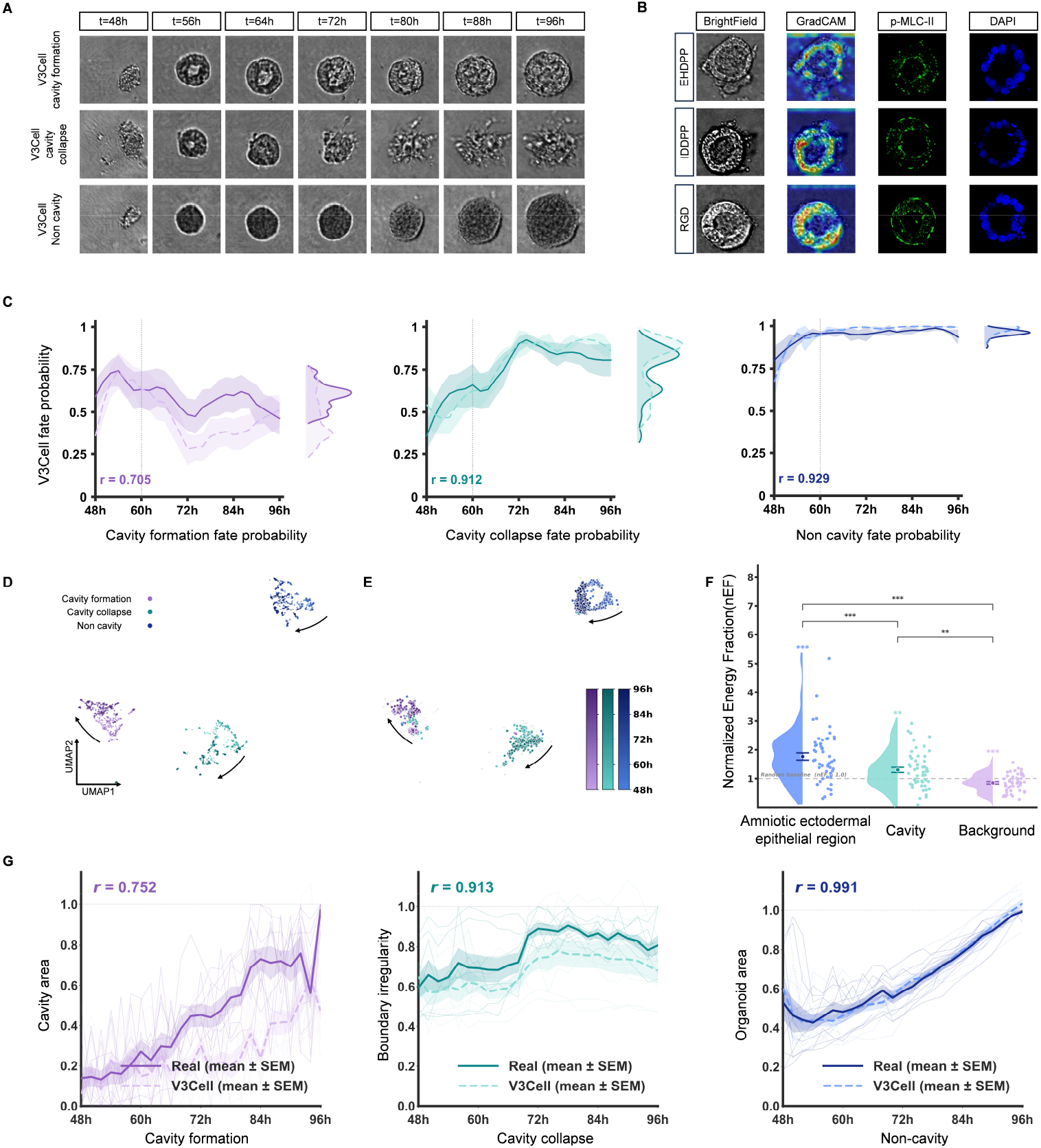
V3Cell predicts organoid fate trajectories and perturbation responses from sparse early frames. All trajectories span culture time 48–96 h. **(A)** Representative V3Cell-constructed virtual fate trajectories for the three fate classes (cavity formation, cavity collapse, and non-cavity). All displayed frames are V3Cell-modeled virtual organoids. **(B)** Grad-CAM spatial attribution and immunofluorescence validation across three perturbation conditions (EHDPP, IDDPP, and RGD). Columns show the brightfield frame, the Grad-CAM map for the cavity-formation logit, phosphorylated myosin light chain II (p-MLC-II), and DAPI. Scale bar, 20 *µ*m for immunofluorescence panels. Brightfield and Grad-CAM panels were cropped to individual organoids. **(C)** Per-frame fate-class probability for real (solid) and virtual (dashed) trajectories in each fate class. Shaded bands denote *±* s.e.m., and the dotted line marks the conditioning boundary. Right marginals show fate-probability distributions as kernel density estimates. *r*, Pearson correlation between real and virtual mean trajectories. **(D)** Unsupervised UMAP projection of held-out real frames using a reducer fitted on real training-frame features only. Points are colored by fate class after projection, and arrows indicate temporal progression. **(E)** The same fixed embedding showing held-out real frames (gray) and the corresponding virtual frames (white-outlined), projected without refitting the reducer or using fate labels. **(F)** Normalized Energy Fraction (nEF) of Grad-CAM attribution across three regions (amniotic ectodermal epithelial region, cavity, and background). The dashed line marks the uniform baseline (nEF = 1). ^*\*\**^*p <* 0.01, ^*\*\*\**^*p <* 0.001 (Wilcoxon signed-rank tests). **(G)** Fate-specific morphometric trajectories (mean *±* s.e.m., thin lines for individual organoids): cavity area for cavity-formation, boundary irregularity for cavity-collapse, and organoid area for non-cavity. Per-class Pearson *r* between real and virtual mean trajectories is shown.

As shown in Fig. 5A (complete sequences in Supplementary Fig. S3), V3Cell produced fate-specific morphological sequences for all three fate classes. Across the displayed culture-time window, the V3Cell-modeled trajectories progressively manifested the hallmark morphological features of their respective fates, yielding visually distinct outputs consistent with the corresponding developmental outcomes.

To test whether fate is decodable from sparse early frames, we applied the fate classifier to the *n* = 36 held-out organoids at the cross-validation-selected early frame (*k*^*\**^ = 4, the fifth conditioning frame). The classifier reached 88.9% accuracy and a macro-*F*_1_ of 0.888 on this held-out set, with performance across all six early frames reported in Supplementary Table S2. The predicted fate was then supplied as the class condition to the frozen image generator that renders each predicted frame.

To quantify temporal fidelity, we compared the per-frame fate-class probability of V3Cell-constructed virtual and matched real trajectories, scored for both by the same independently trained classifier (Fig. 5C). The virtual mean trajectories tracked the real ones in both temporal profile and absolute level, with strong agreement for non-cavity and cavity collapse (Pearson *r* = 0.929 and 0.912) and moderate agreement for cavity formation (*r* = 0.705), consistent with the higher intrinsic variability of the cavity-formation fate. The virtual and real fate-probability distributions also overlapped closely (right marginals, Fig. 5C), supporting that V3Cell constructs fate-consistent morphological trajectories from sparse early frames.

To visualize trajectory organization without imposing fate labels on the embedding, we fitted an unsupervised UMAP reducer exclusively on L2-normalized features extracted from real training frames using an independent ImageNet-pretrained ResNet-50. Held-out real frames were projected into the fixed embedding without refitting. V3Cell-generated frames were then projected using the same reducer, again without refitting or label supervision. The generated frames occupied regions overlapping with the corresponding held-out real trajectories and exhibited similar temporal progression (Fig. 5D,E).

We next asked whether the regions driving V3Cell’s fate encoding correspond to biologically meaningful structures. Grad-CAM attribution for the cavity-formation logit concentrated in the amniotic ectodermal epithelial region rather than in the cavity or background, and this preferential concentration was quantified by the Normalized Energy Fraction (nEF), which was significantly above the uniform baseline in the epithelial region and below it in the background and cavity (Fig. 5F). The attended epithelial region spatially coincided with p-MLC-II immunofluorescence, a marker of actomyosin contractility, and this spatial pattern was consistent across the EHDPP, IDDPP, and RGD perturbation conditions (Fig. 5B), indicating that model attribution localizes to mechanically active epithelial regions in an anatomically grounded and perturbation-generalizable manner.

Finally, we assessed whether V3Cell reproduces quantitative morphodynamics by comparing fate-specific morphometric features between real and virtual sequences over the full time series (Fig. 5G). Real and V3Cell mean trajectories were closely aligned for cavity area in cavity-formation (*r* = 0.752), boundary irregularity in cavity-collapse (*r* = 0.913), and organoid area in non-cavity (*r* = 0.991), supporting that V3Cell captures fate-specific morphological dynamics rather than merely class-level appearance.

## 3 Discussion

The virtual cell concept has emerged as a promising paradigm for modeling cellular behavior in silico. However, existing implementations predominantly operate in molecular feature space at the single-cell level, producing static endpoint predictions from destructive assays. V3Cell extends this paradigm to the organoid scale by constructing virtual 3D cells directly from non-invasive brightfield microscopy. Unlike molecular-space models that require cell lysis for transcriptomic readout or image-based models that depend on fluorescent staining, V3Cell derives all representations from bright-field images alone. This non-destructive approach preserves the intact 3D architecture of the specimen and enables longitudinal tracking of the same organoid over extended culture periods. Our results demonstrate that this modality encodes sufficient morphological information to support both high-fidelity static construction and temporally faithful trajectory prediction.

Our multi-dimensional evaluation demonstrates that virtual 3D cells are not merely visually plausible but closely aligned with real organoids at both the population and single-ROI levels. While distributional fidelity confirmed by IS and KID captures population-level alignment, the Haralick texture analysis and lineage-specific morphometrics establish that virtual 3D cells preserve the fine-grained biological features that define organoid identity. The practical value of this fidelity is evidenced by the phenotyping experiments, where augmenting sparse physical samples with virtual 3D cells improved classification performance by up to 44.7 percentage points and reduced inter-run variability from 15.9% to 1.3%, addressing a practical bottleneck in organoid-based screening where physical sample availability is often limited.

The longitudinal fate prediction results reveal that developmental fate information is already encoded in early-frame brightfield morphology. By first predicting the developmental fate from sparse early observations and then conditioning trajectory generation on this prediction, V3Cell factorizes the task into two well-defined subproblems: inferring the discrete developmental outcome and modeling the corresponding spatiotemporal trajectory. This explicit factorization makes the predictions structured and interpretable. The strong concordance between predicted and real fate-probability dynamics across all three fate classes suggests that V3Cell captures not only the end-point morphology but also the temporal progression toward each fate. Importantly, both fate prediction and trajectory modeling are driven entirely by visual features from early brightfield frames, requiring no knowledge of the applied perturbation at inference time. This property is particularly relevant for screening applications, where the goal is to assess compound effects from observable phenotypic changes without prior knowledge of the mechanism of action.

This result clarifies why V3Cell is formulated as a phenotype-conditioned rather than perturbation-conditioned framework. Many perturbation-response models extrapolate from observed to unobserved perturbations by learning a transferable response structure in molecular or latent feature space, as illustrated by latent-state arithmetic in scGen [14], compositional perturbation embeddings in CPA [15], and gene-regulatory graph embeddings in GEARS [16]. Such extrapolation is most reliable when perturbations occupy a representation in which related inputs induce related responses. In the OPFR panel examined here, however, molecular structure did not provide such a predictive coordinate system: neither structure-derived descriptors nor learned chemical embeddings separated toxic from non-toxic compounds, and structure-based prediction of the developmental readout was no better than chance at screening-relevant accuracy (Supplementary Figs. S4 and S5). This observation is consistent with recent evidence that perturbation-effect predictors can fail to out-perform simple baselines under distribution shift [27]. It also reflects the phenotype being modeled: cavity formation is an emergent tissue-level program, and structurally related analogues can produce divergent developmental outcomes. V3Cell therefore avoids requiring a perturbation-feature prior. By conditioning on early brightfield morphology rather than perturbation identity, it asks whether the effect of an unknown perturbation has already become observable in the developing organoid and uses that observed phenotypic state to forecast future fate. Thus, V3Cell generalizes from sparse phenotypic observation to future trajectory, rather than from chemical structure to response.

The foreground-aware training strategy addresses a practical challenge specific to organoid imaging. Unlike single-cell microscopy, where individual cells can be cleanly segmented, organoid imaging fields contain substantial background artifacts including Matrigel, debris, and out-of-focus structures. By spatially reweighting gradient penalties toward the organoid foreground, V3Cell learns to prioritize biologically meaningful features while suppressing these confounds, as confirmed by the faithful recapitulation of organoid-centric morphometrics such as ellipticity. We anticipate that this vision-guided paradigm will complement existing molecular-space approaches, providing a scalable and longitudinally compatible tool for organoid biology and drug discovery.

### 3.1 Limitations of the study

V3Cell currently models three-dimensional organoids from two-dimensional bright-field projections. Although this imaging strategy enables non-invasive longitudinal observation, it does not directly provide depth-resolved information about features such as internal cavity topology or layered cellular organization. Extending V3Cell to light-sheet microscopy or other three-dimensional imaging modalities could support volumetric modeling when such data are available. In addition, the temporal framework was evaluated in one hESC-derived amniotic-sac organoid system. Future studies across additional organoid systems, cell lines, genetic perturbations and combinatorial treatments will be needed to establish its broader applicability. Finally, the present study focuses on phenotype-conditioned forecasting rather than molecular mechanism inference. Paired brightfield and molecular measurements could complement the current non-invasive framework by linking predicted morphological trajectories to their associated molecular processes.

## 4 Methods

### 4.1 Datasets

Two independent imaging datasets were used to support the two stages of the V3Cell framework. The first, a static brightfield image dataset, comprises three organoid lineages (colon, stomach, and lung) assembled from publicly accessible repositories. Raw microscopy fields for the colon, stomach, and lung lineages were obtained from the OrgaExtractor repository [68], the in-house collection released alongside our group’s previous work [69], and the Orga-Dete lung organoid dataset [70], respectively. No pre-cropped ROIs or pre-existing phenotype labels from the original sources were used. Individual organoid ROIs were cropped in-house from the raw fields. The resulting dataset comprises 17,331 ROI images distributed across 9 morphological phenotype classes, including 4 colon classes (2,476 images), 3 stomach classes (897 images), and 2 lung classes (13,958 images). For each ROI, a binary foreground mask delineating the organoid boundary from the culture background was derived from an organoid instance segmentation model developed in our group [71]. Each ROI image served as a real training target for Stage 1, and its corresponding mask was used only to define training-time foreground constraints. Morphological phenotype labels and foreground-mask verification were produced under the annotation workflow described below.

The second dataset is a longitudinal brightfield time-lapse dataset established to capture the developmental trajectories of living hESC-derived amniotic-sac organoids (rosette-like 3D structures) under chemical perturbation. It is biologically and experimentally distinct from the static dataset described above and was collected specifically to support temporal modeling in the second stage. The human embryonic stem cell (hESC) line H1 was obtained from the Shao laboratory (School of Aerospace Engineering, Tsinghua University) and maintained on Matrigel-coated plates in mTeSR1 medium (85850, STEMCELL Technologies). Cells were seeded as single cells at 2 × 10^4^ cells cm^−2^ onto a thick Matrigel bed with nominal thickness ≥ 100 *µ*m. Immediately after passaging, the culture medium was supplemented with 10 *µ*M Y-27632 (1254/10, Tocris). After 24 h, the medium was replenished with fresh mTeSR1 containing 4% (v/v) Geltrex (A1413201, Thermo Fisher Scientific) and replenished daily thereafter. On day 2 of culture, cells were treated with 2-ethylhexyl diphenyl phosphate (EHDPP) at a final concentration of 10 *µ*M (treatment group) or an equivalent volume of DMSO (vehicle control). To further probe the generalizability of model attention beyond a single compound, two additional perturbation conditions were acquired under the identical culture and imaging protocol: isodecyl diphenyl phosphate (IDDPP), a second organophosphate flame retardant, at a final concentration of 10 *µ*M, and the RGD peptide, a positive-control integrin *β*_1_ antagonist, at 1 *µ*M. The IDDPP and RGD single-organoid time-lapse sequences (*n* = 10 and *n* = 10, respectively) were reserved exclusively for the Grad-CAM attention-generalization analysis described in the Grad-CAM spatial attribution and normalized energy fraction sub-section and were not used to train or test the temporal model. Continuous brightfield time-lapse imaging was performed on a DeltaVision Elite microscope (Applied Precision) equipped with a 10× objective, acquiring one frame every 15 min over a 2-day window, within a stage-top live-cell incubator maintained at 37^*°*^C and 5% CO_2_. Each time-lapse sequence was assigned to one of three developmental fate categories. The *cavity-formation* phenotype was characterized by cells arranging around a shared point of apical constriction before approximately 64 h, followed by the emergence of a rosette-like structure surrounding a central cavity between 72 and 96 h. The *non-cavity* phenotype was characterized by the absence of such a structure throughout the imaging window. The *cavity-collapse* phenotype was characterized by a malformed rosette in which the central cavity failed to stabilize. The three fate outcomes were observed in both treatment and control groups but occurred with substantially higher frequency under EHDPP perturbation, indicating a probabilistic rather than deterministic perturbation response. The training corpus for the temporal model comprises 245 labeled time-lapse sequences (28 cavity-formation, 106 cavity-collapse, and 111 non-cavity), and an additional class-stratified set of 36 sequences (12 per fate class), disjoint from these 245, was reserved as a held-out test set for evaluation. Fate-label assignment followed the annotation workflow described below. Each raw sequence contained 196 frames acquired at 15-min intervals over the 48-h observation window from culture time 48 to 96 h, with chemical treatment administered at 48 h. For temporal modeling, each sequence was downsampled by retaining every eighth frame, yielding 25 frames at 2-h intervals.

Static images used for the reported Stage 1 experiments were resized to 64 × 64 pixels. The generator architecture supports configurable output resolutions. For the reported temporal experiments, the amniotic-sac generator and generated frames were trained and evaluated at 64 × 64 pixels to reduce computational cost. For downstream morphometric segmentation, real and generated frames were resized to 256 × 256 pixels using bilinear interpolation. Pixel intensities were linearly normalized to [ −1, 1], class labels were encoded as one-hot vectors **c** ∈ {0, 1} ^C^, and horizontal flipping was applied as data augmentation during all training stages.

### 4.2 Annotation workflow and quality control

All phenotype and fate labels used in V3Cell were produced in-house under a unified annotation workflow designed to ensure consistency across lineages and between the static and dynamic datasets. Raw imaging data were first pre-processed and deidentified, after which three trained laboratory researchers independently performed annotation using a shared labeling schema. No pre-existing labels from the public source repositories were used.

For the static brightfield dataset, morphological phenotype labels were assigned per ROI following the phenotype taxonomy and labeling protocol established in our previous work [69, 72], which defines 4 colon, 3 stomach, and 2 lung classes on the basis of reproducible morphological criteria (e.g., budding status, lumenization, overall shape regularity). Each ROI was annotated independently by three researchers. The consensus label (agreement of at least two annotators) was adopted as the final class label, and ROIs with no majority agreement were resolved by a senior expert reviewer. In parallel, each foreground mask produced by the OrgLine [71] instance segmentation model was visually inspected frame-by-frame. Masks with leakage, under-segmentation, or mis-attribution to neighboring organoids were manually corrected using a polygon-editing tool, and the corrected masks were cross-checked by a second annotator before entering the training set.

For the longitudinal time-lapse dataset, each sequence was annotated at the sequence level using the three fate categories defined in the Datasets subsection (cavity-formation, non-cavity, cavity-collapse). Annotation was performed by reviewing the full 2-day recording with specific attention to the 64–96 h window, during which fate-diagnostic rosette dynamics emerge. Three trained researchers labeled every sequence independently while remaining blinded to the experimental condition (EHDPP treatment or DMSO control) to prevent condition-driven labeling bias. The consensus label was adopted when at least two annotators agreed, and ambiguous cases—in particular borderline cavity-collapse sequences where the central cavity partially stabilized— were resolved by expert consensus involving a senior domain expert. Inter-annotator agreement was monitored on a held-out calibration subset prior to full-scale annotation to confirm that the three annotators achieved substantial concordance on the fate taxonomy.

### 4.3 Overview of the V3Cell framework

V3Cell is a two-stage framework that formalizes virtual 3D cell modeling as two complementary conditional generation tasks. In the static task, given a one-hot class label **c** ∈ {0, 1} ^*C*^, a conditional model *p*_*θ*_(**x** | **c**) is trained such that each sample 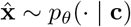 constitutes a static virtual 3D cell. In the dynamic task, given *K* = 6 early brightfield frames of a live organoid, a fate classifier Φ first predicts the developmental outcome *ĉ* = Φ(**x**_*k*_^*\**^) from a single early frame. Here, *k*^*\**^ = arg max_*k*_ Acc(Φ(**x**_*k*_), *c*) is the most informative frame index selected by 5-fold cross-validation (*k*^*\**^ = 4). The dynamic model then produces the complete predicted trajectory 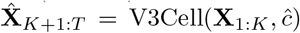 from the same *K* = 6 conditioning frames. Neither stage requires knowledge of the applied perturbation at inference time. Both are driven entirely by morphological features extracted from the brightfield observations. An overview of the complete architecture is shown in Figure 6.

**Fig. 6.**
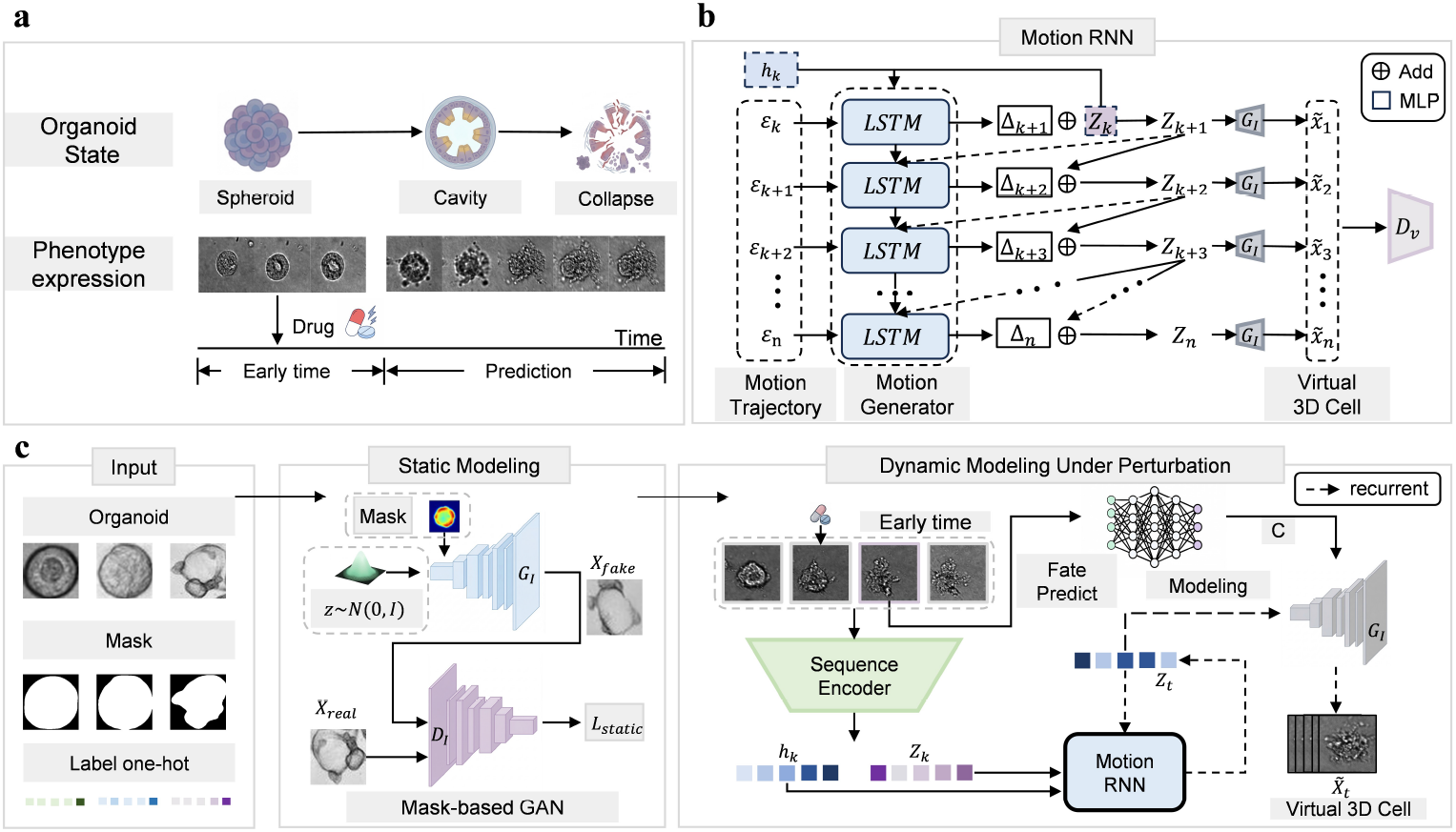
Overview of the V3Cell framework for virtual 3D cell modeling. **(a)** Conceptual illustration of the virtual 3D cell paradigm. Organoid phenotypic state is modeled as a trajectory in a continuous latent space, progressing from early developmental states (spheroid) through fate-specific outcomes (cavity formation or collapse). External chemical perturbations shift the developmental trajectory, enabling fate prediction and fate-conditioned morphological modeling from sparse early-frame observations. **(b)** Architecture of the Stage 2 Motion RNN. The module autoregressively evolves a motion latent by combining the previous motion state with stochastic noise inputs through a multi-layer LSTM, producing incremental latent updates that are decoded by the frozen Image Generator to render predicted future frames 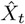 . **(c)** End-to-end pipeline of the V3Cell framework. Left (Stage 1, foreground-aware static virtual 3D cell construction): a class-conditioned GAN trained with foreground-weighted constraints maps latent noise *z* ∼ *N* (**0, I**) and one-hot phenotype label *c* to high-fidelity brightfield images of static virtual 3D cells. Foreground masks are used only during training to weight foreground-focused losses and are not supplied at inference. Right (Stage 2, dynamic virtual 3D cell modeling and fate prediction): early-frame brightfield observations are encoded by a Sequence Encoder into content and motion latents, a parallel Fate Prediction module identifies the developmental outcome *ĉ* from an optimal observation frame 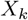, and MotionRNN uses *ĉ* as its categorical conditioning signal while evolving the motion latent. The combined latent representation is decoded by the frozen Image Generator to produce the complete predicted morphological trajectory of the virtual 3D cell.

### 4.4 Foreground-aware static virtual 3D cell generator

Static virtual 3D cells were modeled by a class-conditional GAN built on the R3GAN framework [73]. For the Stage 1 static generators, the generator *G* maps a latent code **z** ∼ *N* (**0, I**) (*d*_*z*_ = 64) and a one-hot phenotype label **c** ∈ {0, 1} ^*C*^ to a brightfield image through a generative basis layer followed by grouped-convolution residual blocks with bilinear upsamplers. The discriminator *D* mirrors this structure. Class conditioning was applied symmetrically to both *G* and *D* in Stage 1. The same decoder backbone was used to train a separate amniotic-sac generator *G*_AS_ for Stage 2, which was then frozen as the image decoder for temporal modeling.

To prioritize biologically meaningful features and suppress background artifacts such as Matrigel and debris, the zero-centered gradient penalties were spatially re-weighted by a foreground-emphasis map:

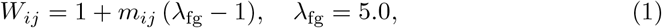

where **m** ∈ {0, 1} ^*H×W*^ is the binary foreground mask used only for training-time loss weighting and not as a generator input at inference. The foreground-weighted R1 and R2 gradient penalties, applied to real and generated samples respectively, were defined as:

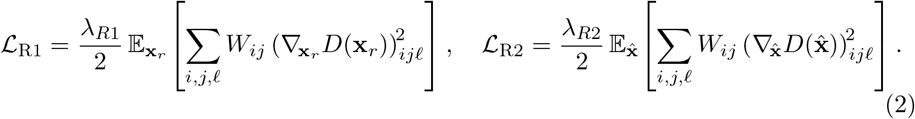

The discriminator and generator were optimized using a relativistic pairing objective with softplus loss:

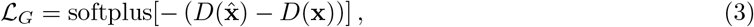

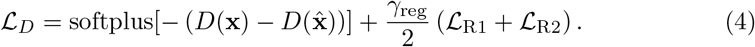

The complete adversarial objective combined full-image and foreground-only branches:

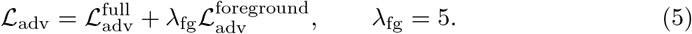

Optimization, scheduling, and inference-time behavior for Stage 1 are described in the Details of pipeline training and inference subsection below.

### 4.5 Dynamic virtual 3D cell modeler

The dynamic modeler was trained on top of a separate frozen amniotic-sac generator *G*_AS_ and comprises three components that operate jointly at inference time: a fate classifier that predicts the developmental outcome from a single early frame **x**_*k*_^*\**^, a sequence encoder that compresses each of the *K* = 6 conditioning frames into a latent representation, and MotionRNN, which autoregressively evolves the latent to generate future frames.

The fate classifier is a ResNet-50 [74] backbone pre-trained on ImageNet [75] with a three-way linear classification head, fine-tuned end-to-end with cross-entropy loss. The optimal observation frame *k*^*\**^ was identified by 5-fold cross-validation across candidate frame indices, with *k*^*\**^ selected as the index that maximized cross-validated classification accuracy. This procedure selected *k*^*\**^ = 4, which was used for held-out fate prediction. At inference time, the predicted fate *ĉ* = Φ(**x**_*k*_^*\**^) ∈ {cavity-formation, non-cavity, cavity-collapse} was used as the class-conditioning input to the frozen generator *G*_AS_ and broadcast across all decoded frames of the trajectory, exactly as in Stage 1 static generation. MotionRNN did not receive *ĉ*; it evolved the motion latent solely from the early-frame encoding and the injected per-step noise.

The sequence encoder *f*_*ϕ*_ processed each early frame through a 7 × 7 convolutional stem, five residual downsampling blocks, global average pooling, and parallel linear heads, yielding a VAE-style reparameterized latent:

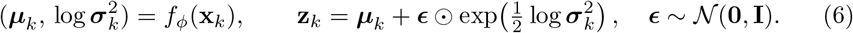

The sequence {**z**_1_, …, **z**_*K*_} was passed through a two-layer LSTM with hidden dimension *d*_*h*_ = 384 to initialize the motion predictor. Each latent was further factorized into a content component **z**_*C*_ ∈ ℝ^32^ (fixed from **z**_1_) preserving organoid-specific appearance, and a motion component **z**_*M*_ ∈ ℝ^32^ carrying temporally evolving dynamics. The motion component was autoregressively evolved by MotionRNN *R*_*ψ*_, which consumed the previous motion state and a stochastic noise input to produce an incremental update:

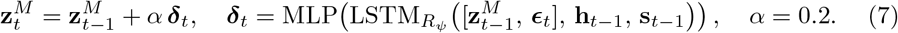

The full latent 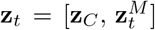 was decoded by the frozen amniotic-sac generator as 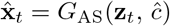, with the predicted fate *ĉ* supplied as the class condition and broadcast to every frame, to produce each predicted future frame. Fate therefore modulates the rendered appearance of each frame, whereas the latent motion dynamics produced by MotionRNN are independent of *ĉ*.

The composite training loss combines reconstruction, adversarial, feature-matching, contrastive, and regularization terms:

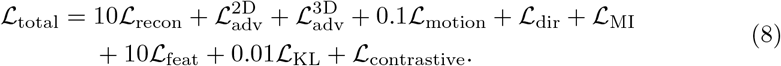

where 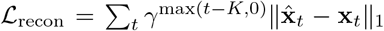 with *γ* = 0.95 is a temporally discounted pixel-wise reconstruction loss. The adversarial terms used a 2D image discriminator and a multi-scale 3D video discriminator, both optimized with relativistic average least-squares GAN objectives. No WGAN-GP term was used in Stage 2. *L*_dir_ enforced biologically consistent morphological progression direction. *L*_MI_ prevented motion mode collapse by maximizing alignment between injected and reconstructed noise [76]. *L*_motion_ penalized abrupt latent transitions, and *L*_KL_ regularized the encoder posterior toward the standard Gaussian prior. Individual loss terms are defined in Supplementary Note 1. Optimization, class-balanced sampling, and inference-time behavior for Stage 2 are described in the Details of pipeline training and inference subsection below.

### 4.6 Details of pipeline training and inference

The two stages of V3Cell were trained sequentially. Three lineage-specific static generators were trained for the colon, stomach, and lung analyses. A separate amniotic-sac generator *G*_AS_ was trained for temporal modeling and then frozen to serve as the fixed decoder during Stage 2 training. Within Stage 2, the fate classifier, sequence encoder, and MotionRNN were optimized on the time-lapse dataset, while gradient flow into *G*_AS_ was disabled.

#### Static generator training and inference

The static generator was trained independently for each lineage (colon, stomach, lung) using the foreground-weighted relativistic pairing objective described above. Optimization used the Adam optimizer with *β*_1_ = 0 and *β*_2_ linearly scheduled from 0.9 to 0.99. The learning rate followed a cosine schedule from 2 × 10^−4^ towards a nominal floor of 5 × 10^−5^ over a 20,000-kimg horizon. Each colon-, lung- and stomach-organoid generator was trained for 443 kimg with a batch size of 64 on a single NVIDIA A100 GPU. The learning rate was 1.998 × 10^−4^ at the selected 443-kimg checkpoint, which was used for all subsequent analyses. The gradient-regularization coefficient followed the same nominal cosine schedule from 150 towards 15. Gradient penalties *L*_R1_ and *L*_R2_ were applied at every discriminator update with the foreground-emphasis weight *λ*_fg_ = 5.0. Before Stage 1 training, 10% of the real images from each lineage were reserved as a fixed stratified test set using random seed 42. These images and their corresponding masks were excluded from generator training and model selection. The remaining 90% of ROI–mask pairs were used to train the lineage-specific generators. At inference, static virtual 3D cells were generated by sampling **z** ∼ *N*(**0, I**) with a fixed target class label **c**. No stochastic augmentation, test-time refinement, or foreground mask was used at inference.

#### Dynamic modeler training and inference

The amniotic-sac generator *G*_AS_ and the temporal model were trained exclusively using the 245 training sequences. All frames from the 36 held-out test sequences were excluded from training and model selection for both components. The fate classifier was fine-tuned on the time-lapse dataset with cross-entropy loss using the Adam optimizer at a learning rate of 1 × 10^−4^ under cosine annealing, for 50 epochs with a batch size of 32. The sequence encoder and MotionRNN were jointly optimized against the composite loss *L*_total_ defined above using the Adam optimizer with *β*_1_ = 0.5, *β*_2_ = 0.999, and a learning rate of 2 × 10^−4^. Each training iteration used *T* = 25 frames with *K* = 6 conditioning frames and a batch size of 32. For temporal training, kimg denotes the cumulative number of processed frames rather than the number of videos, kimg = Σ _steps_(*B* × *T*)*/*1000. Optimization proceeded for 581 kimg on 8 NVIDIA A100 GPUs. The learning rate was held fixed at 2 × 10^−4^ throughout Stage 2 training, with no learning-rate schedule. Class imbalance across the three fate categories (28 cavity-formation, 106 cavity-collapse, and 111 non-cavity sequences) was addressed by class-balanced sampling, ensuring that each mini-batch contained approximately equal proportions of all three fate classes. Throughout Stage 2 training, *G*_AS_ remained frozen. At inference, given the *K* = 6 early frames of an unseen organoid, the sequence encoder produced the initial latent factorization into content and motion components. The fate classifier read the cross-validated frame **x**_*k*_^*\**^ (*k*^*\**^ = 4) to emit *ĉ*. Motion-RNN autoregressively rolled out the motion latent for the remaining *T* − *K* = 19 frames, which were decoded by *G*_AS_ to yield the complete 25-frame predicted trajectory 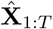 . No ground-truth future frames, foreground masks, or perturbation identity were provided at inference time.

### 4.7 Grad-CAM spatial attribution and normalized energy fraction

To assess whether V3Cell encodes fate-relevant information from spatially specific anatomical regions, we applied gradient-weighted class activation mapping (Grad-CAM) [77] to the V3Cell image encoder. Because the encoder outputs a 64-dimensional latent vector **z** rather than class logits, we trained a linear fate probe on frozen encoder representations to provide class-discriminative gradients for post hoc attribution. The probe was trained on the EHDPP dataset comprising 245 videos across three fate classes: cavity-formation (*n* = 28), cavity-collapse (*n* = 106), and non-cavity (*n* = 111). It consisted of a single fully connected layer optimized with Adam for 200 epochs. Its training accuracy was monitored only to confirm that the frozen encoder representations retained fate-related information. The probe was used solely to provide class-discriminative gradients for post hoc attribution and was never used to optimize generated frames during inference. The Grad-CAM target layer was the encoder’s third convolutional block, which retained a 32 × 32 spatial resolution.

Grad-CAM and nEF analyses were restricted to videos exhibiting cavity formation or a cavity-forming tendency, so as to align the attribution analysis with the immunofluorescence validation described below. The analyzed subset comprised 58 videos: 28 EHDPP training videos, 10 EHDPP held-out videos, 10 IDDPP videos, and 10 RGD videos. Because this analysis aimed to localize fate-relevant encoder responses rather than to estimate predictive generalization, both training and held-out EHDPP cavity-forming videos were included.

For each frame, the cavity-formation logit *s*_cavity_ from the linear probe was back-propagated to the target convolutional layer. Let *A*^*k*^ ∈ ℝ^H*×*W^ denote the *k*-th activation map of the target layer. The Grad-CAM channel weight was computed as

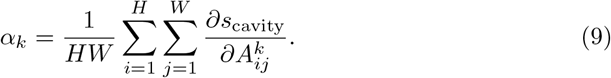

The frame-level Grad-CAM activation map was then computed as

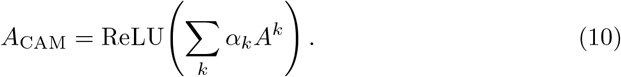

The resulting map was normalized to [0, 1] and upsampled to the original frame resolution of 256 × 256 pixels by bilinear interpolation.

For post hoc attribution quantification, spatial regions were defined from per-frame segmentation masks produced by task-specific OrgLine [71] segmentation models. A whole-organoid boundary model produced the foreground mask, and a cavity model produced the cavity mask. All masks were manually inspected and corrected where necessary prior to analysis. Three mutually exclusive regions were defined: background (pixels outside the whole-organoid mask), cavity (pixels inside the cavity mask), and epithelial cell ring (pixels inside the whole-organoid mask but outside the cavity mask).

These segmentation masks were used only for post hoc regional quantification and were not provided to V3Cell during inference.

For each region ℛ in each frame, we first computed the Energy Fraction (EF), the fraction of total Grad-CAM activation falling inside that region:

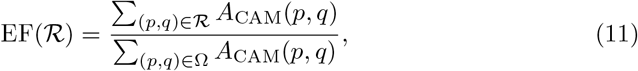

where Ω is the full image domain. To account for differences in region size, EF was normalized by the area fraction of the corresponding region:

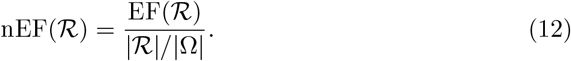

An nEF of 1 indicates Grad-CAM activation proportional to region area, corresponding to a spatially uniform attention baseline. Values greater than 1 indicate preferential attention to a region, whereas values below 1 indicate relative suppression.

Frames with zero total Grad-CAM activation were excluded from the nEF computation. nEF was computed for every remaining frame of each analyzed video and averaged across frames to obtain one video-level nEF per region. Statistical analyses were performed at the video level on these per-video mean nEF values: one-sample Wilcoxon signed-rank tests compared each region against the uniform baseline of nEF = 1, and paired Wilcoxon signed-rank tests were used for between-region comparisons.

### 4.8 Immunofluorescence validation of model attribution

To assess whether the regions highlighted by V3Cell Grad-CAM attribution correspond to biologically relevant anatomical structures, we compared the attribution maps with immunofluorescence (IF) staining from the same amniotic-sac organoid experimental system. IF images were obtained from the dataset provider and followed the previously reported staining protocol [78]. These fluorescence images were used only for post hoc anatomical validation of model attribution and were not used for V3Cell training, fate prediction, trajectory generation, or model selection.

Organoids were fixed in 4% paraformaldehyde for 60 min at room temperature, permeabilized with 0.25% Triton X-100 for 30 min, and blocked in 3% bovine serum albumin (BSA) for 60 min at 37^*°*^C. Samples were incubated with primary antibodies overnight at 4^*°*^C, followed by fluorophore-conjugated secondary antibodies for 60 min at room temperature, and nuclei were counterstained with DAPI (1:5000, Sigma, D9542) for 30 min. Fixed samples were imaged using a high-speed spinning-disk confocal microscope (Andor Technology) equipped with a Leica 0.45 NA 10× air objective. Primary antibody sources, catalog numbers, and working dilutions followed the dataset-provider protocol and are summarized in Table S3.

We focused on phosphorylated myosin light chain II (p-MLC-II), a marker of actomyosin contractility, because cavity formation in this amniotic-sac organoid system involves epithelial organization and actomyosin activity around the nascent cavity.

Developmental marker channels, including OCT4, TFAP2A, CDX2, and DAPI, were used to provide anatomical and cellular context, whereas p-MLC-II served as the primary IF marker for interpreting whether model attribution localized to mechanically active epithelial regions.

Attribution maps and region-level Normalized Energy Fraction (nEF) were computed as described in the Grad-CAM spatial attribution and normalized energy fraction subsection, using the same three anatomical regions: non-organoid background, cavity, and epithelial cell ring. IF images were used to interpret the anatomical relevance of regions with elevated Grad-CAM nEF. In particular, enrichment of attribution in the epithelial cell ring was compared with the spatial localization of p-MLC-II in representative matched or condition-matched IF fields. Because IF imaging required fixation, fluorescence images were used as endpoint or representative anatomical validation rather than as longitudinal inputs to V3Cell.

### 4.9 Evaluation metrics

#### Kernel Inception Distance (KID)

Given *m* real-image feature vectors 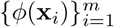 and *n* virtual-image feature vectors 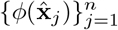 extracted by an ImageNet-pretrained Inception-v3 feature extractor *ϕ*(·), KID was computed as the squared maximum mean discrepancy (MMD) under a degree-3 polynomial kernel 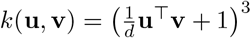 :

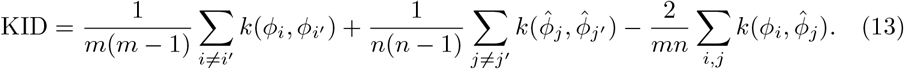

A subtype-balanced sampling protocol with *n* = min_*s*_ min(*N*_real,*s*_, 2000) was used to remove subtype-size confounds, with subtype-balanced samples merged across subtypes, and KID was averaged over 10 independent sampling runs.

#### Inception Score (IS)

Let 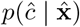 denote the softmax output of the ImageNet-pretrained Inception-v3 classifier evaluated on virtual image 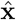, and 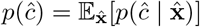 the corresponding marginal. The Inception Score was computed as:

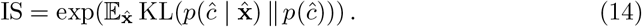

Following standard practice, IS was computed over all constructed images across 10 independent runs and reported as mean ± standard deviation across runs.

#### Haralick texture descriptors

For each ROI, a gray-level co-occurrence matrix (GLCM) was computed from the 8-bit grayscale image restricted to the foreground mask. For a displacement vector (Δ*x*, Δ*y*), each entry of the unnormalized GLCM counts co-occurring intensity pairs:

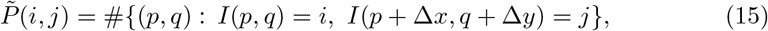

and is normalized to 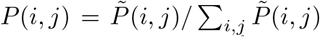 . GLCMs were computed at four distances {1, 2, 3, 4} pixels and four angles 0^*°*^, {45^*°*^, 90^*°*^, 135^*°*^} at 256 gray levels and averaged across angles for rotational invariance. Five Haralick descriptors [65] were then derived:

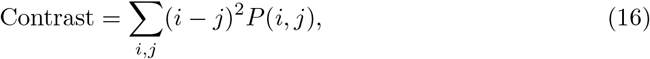

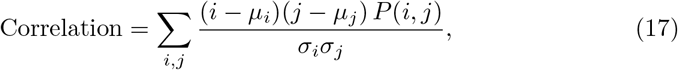

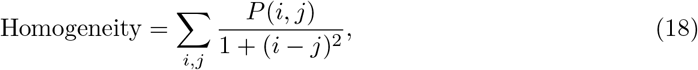

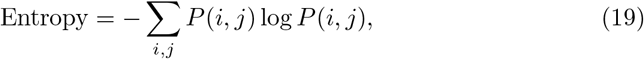

where *µ*_*i*_, *µ*_*j*_ and *σ*_*i*_, *σ*_*j*_ are the means and standard deviations of the marginal distributions *P*_*i*_(*i*) = Σ_*j*_ *P* (*i, j*) and *P*_*j*_(*j*) = Σ_*i*_ *P* (*i, j*). The Information Measure of Correlation 1 (IMC1) was computed as:

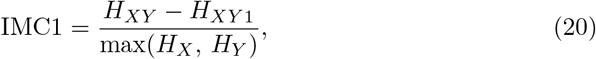

where *H*_*X*_ = − Σ_*i*_ *P*_*i*_(*i*) log *P*_*i*_(*i*), *H*_*Y*_ = − Σ_j_ *P*_*j*_(*j*) log *P*_*j*_(*j*), *H*_*XY*_ = *− Σ*_*i*_ *P*_*i*_ (*i*) log *P*_*i*_ (*i*), *H*_*XY*_ *= − Σ*_*i,j*_ *P* (*i, j*) log *P* (*i, j*) (the joint entropy, equivalent to the Entropy descriptor defined above), and *H*_*XY1*_ = *− Σ*_*i,j*_ *P* (*i, j*) log *P*_*i*_(*i*) *P*_*j*_(*j*) . IMC1 is non-positive by construction and quantifies the degree to which the joint GLCM distribution departs from the product of its marginals.

Real and V3Cell descriptor distributions were compared with two-sided Wilcoxon rank-sum tests, and effect sizes were quantified by Cohen’s *d* [66]:

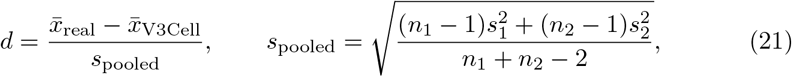

with |*d*| *<* 0.2 regarded as biologically negligible, 0.2 ≤ |*d*| *<* 0.5 as small, and |*d*| ≥ 0.5 as moderate or larger.

#### Lineage-specific morphometric descriptors

Three lineage-specific morphometric descriptors were extracted per ROI:

- *Bud number* (colon): the count of morphologically distinct convex protrusions extending from the organoid boundary, where each bud was defined as a discrete outgrowth with a visible constriction at its base relative to the main organoid body. Bud numbers were assigned by human annotators following the consensus labeling protocol described in the Annotation Workflow subsection above.
- *Lumen area* (lung): normalized internal-hollow-region area,

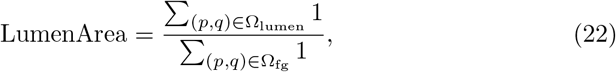

where Ω_fg_ is the foreground mask and Ω_lumen_ ⊂ Ω_fg_ is the internal hollow region detected within it.
- *Ellipticity* (stomach): computed from the major and minor axes *a* and *b* of the best-fitting ellipse of the foreground region,

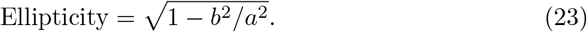

Distributional comparisons between real and virtual populations used the Wilcoxon rank-sum test and Cohen’s *d* as defined above.

#### UMAP visualization

For static phenotypic visualization, a *D*-dimensional feature vector **f**_*i*_ ∈ ℝ^D^ was extracted per sample using ResNet-101 features at *D* = 2048, L2-normalized, and projected into ℝ^2^ via UMAP [67]. The static visualization used `n_neighbors` = 15, `min_dist` = 0.1, cosine metric, and random state = 42.

For trajectory visualization, L2-normalized features extracted using an independent ImageNet-pretrained ResNet-50 were obtained from frames in the 245 training sequences. These features were used to fit UMAP without label supervision (`n_neighbors` = 30, `min_dist` = 0.25, cosine metric and random state = 42). The fitted reducer was subsequently fixed and used, without refitting, to project frames from the 36 held-out real trajectories and their corresponding V3Cell-generated trajectories. Fate labels were used only to color the points after projection.

#### Phenotyping augmentation classifier

To evaluate whether V3Cell-generated virtual organoids improve downstream phenotype classification under sparse-sampling conditions, we trained an independent phenotype classifier for each organ lineage. The classifier used a ResNet-50 backbone initialized with ImageNet-pretrained weights, with the final fully connected layer replaced by a lineage-specific linear classification head mapping the 2048-dimensional feature vector to the number of phenotype classes (colon with 4 classes, stomach with 3 and lung with 2). All layers were fine-tuned end-to-end. Grayscale brightfield ROIs were replicated into three channels, resized to 64 × 64 pixels, and normalized using ImageNet channel statistics.

For each lineage, 10% of real images were held out as a fixed stratified test set (random seed 42). This test set contained only real images and was never used for training, validation, model selection, or virtual-image generation. The remaining real images formed the training pool. To simulate low-resource phenotyping, we sampled *N* ∈ {10, 25, 50} real images per class from the training pool and compared three mutually exclusive training conditions using identical real-image splits: real-only, standard augmentation, and V3Cell augmentation. In the real-only condition, the classifier was trained only on the sampled real images. In the standard-augmentation condition, the same sampled real images were augmented online using random horizontal and vertical flips (*p* = 0.5), random rotations within ± 15^*°*^, and color jitter in brightness, contrast, and saturation (magnitude 0.2). Because each image was randomly transformed on the fly at every access, this condition increased sample diversity without changing the number of training samples, leaving the per-class training cardinality fixed at *N* . In the V3Cell-augmentation condition, the same sampled real images were supplemented with an equal number of class-matched V3Cell-generated virtual ROIs (a 1:1 real-to-virtual ratio, i.e., *N* virtual ROIs per class), added as stored images to the training set. A virtual-only setting with *N* = 0 real images per class was also evaluated to test whether V3Cell-generated images alone contained class-discriminative morphological information.

Synthetic images were added only to the training set and were never included in validation or test sets. Validation images were sampled from the real-image training pool, disjoint from the sampled training images, and used only for model selection. Models were trained with cross-entropy loss and the Adam optimizer (learning rate 1 × 10^−3^, batch size 128) for up to 50 epochs. The learning rate was reduced by a factor of 0.5 when validation accuracy plateaued for 5 epochs, and early stopping was applied if validation accuracy did not improve for 10 consecutive epochs. The checkpoint with the highest validation accuracy was restored for evaluation.

Each configuration was repeated across five random seeds (0–4) using matched real-image splits across conditions. Primary performance was quantified by macro-averaged *F*_1_ on the fixed real test set. The 25-image-per-class setting was used for the main augmentation-effect and reproducibility analyses shown in Fig. 4B–C.

#### Macro-averaged F_1_ for phenotyping augmentation

Phenotyping performance was quantified by the macro-averaged *F*_1_ score. For each phenotype class *c* ∈ {1, …, *C* }, precision and recall were computed from the confusion matrix entries:

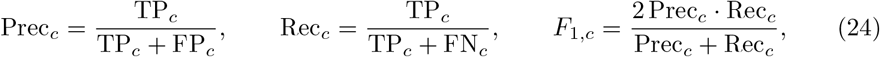

and the macro-averaged score treats every class equally regardless of class frequency:

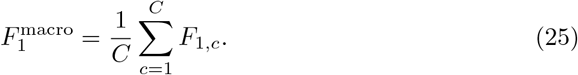

Across seeds, the standard deviation of *F* ^macro^ served as the primary reproducibility metric.

#### Per-frame fate-class probability for trajectory fidelity

Temporal fidelity of V3Cell-constructed virtual trajectories was quantified by the per-frame fate-class probability for the ground-truth fate class. For a frame **x**_*t*_ (real) or 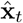 (virtual) and a fixed fate classifier Φ producing logits **z** = Φ( ·) ∈ ℝ^3^, the probability of fate class *c* was defined as the softmax value:

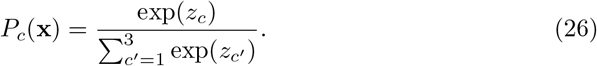

For each test organoid of ground-truth fate *c*^⋆^, the sequence 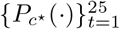 was computed over the 25-frame horizon (*K* = 6 conditioning frames, 19 predicted frames). Mean probability trajectories were then averaged across organoids within each fate class and smoothed by a Savitzky–Golay filter [79] with window length 7 and polynomial order 3 to suppress high-frequency noise. The same procedure was applied to real frames to obtain paired reference curves.

#### Fate-specific morphometric trajectories

Three fate-specific morphological readouts were extracted independently from each frame using ROIRefineNetV1 with an Atto backbone. Real and V3Cell-generated frames were resized to 256 × 256 pixels using bilinear interpolation before segmentation. Segmentation logits were transformed using a sigmoid function and binarized at a threshold of 0.5. Separate task-specific checkpoints were used for cavity-formation, cavity-collapse and non-cavity organoids.

For cavity-formation organoids, cavity area was defined as the number of foreground pixels in the segmented cavity mask. For cavity-collapse organoids, boundary irregularity was calculated from the largest connected component of the segmented organoid outline as *P* ^2^*/*(4*πA*), where *A* and *P* denote area and perimeter, respectively. For non-cavity organoids, organoid area was defined as the number of foreground pixels in the general organoid mask.

For each video, both the real and corresponding generated trajectories were normalized by the peak value of the real trajectory. Mean trajectories were calculated across videos at each time point, and agreement was quantified using Pearson correlation between the real and generated mean trajectories.

### 4.10 Empirical study

To clarify how each reported result in the Results section was obtained, we summarize below the experimental protocol associated with each of the four major analyses. Unless otherwise noted, all analyses used the V3Cell models described above, A100 GPUs, and the dataset splits specified in the Datasets subsection.

#### Experimental setup for static virtual 3D cell generation

This experiment evaluates the ability of Stage 1 to synthesize high-fidelity, class-consistent static virtual 3D cells for each of the nine morphological phenotype classes across the colon, stomach, and lung lineages. From the complete collection of 17,331 static ROI–mask pairs, 10% of images from each lineage were reserved as a fixed stratified test set before generator training. The remaining 90% were used to train the lineage-specific generators. A separate lineage-specific foreground-aware static generator was trained using the Stage 1 protocol specified in the Details of pipeline training and inference subsection. Generation quality was quantified by comparing a class-balanced population of constructed virtual 3D cells against the full set of real ROIs using KID, IS, and qualitative side-by-side inspection. Each generator was sampled to produce a class-balanced virtual population used both for the fidelity evaluations below and for the phenotyping augmentation experiments.

#### Experimental setup for multi-dimensional morphological and texture fidelity evaluation

This experiment evaluates whether V3Cell-constructed static virtual 3D cells reproduce the morphological and textural feature distributions of real organoids at a population level. For each lineage, the full set of real ROIs was compared with an equally sized set of V3Cell-constructed ROIs sampled from the corresponding lineage-specific generator. Haralick texture descriptors (contrast, correlation, homogeneity, entropy, and IMC1) were computed per ROI on the 8-bit grayscale image restricted to the foreground mask, using the GLCM protocol specified under the Haralick texture descriptors subsection. Lineage-specific morphometric descriptors, bud number for colon, lumen area for lung and ellipticity for stomach, were extracted per ROI as specified under the Lineage-specific morphometric descriptors subsection. For each descriptor, real versus V3Cell distributions were compared with a two-sided Wilcoxon rank-sum test, and effect size was quantified by Cohen’s *d*, with | *d*| *<* 0.2 regarded as biologically negligible. UMAP projections of ResNet-101 features were computed with the static-visualization hyperparameters to provide a class-level structural comparison.

#### Experimental setup for phenotyping augmentation

This experiment evaluates whether V3Cell-constructed virtual populations can augment small real training sets for downstream phenotype classification. For each lineage, a ResNet-50 phenotype classifier was fine-tuned under physical-sample budgets of {0, 10, 25, 50} real ROIs per class (with 0 corresponding to the virtual-only setting) and three augmentation conditions (real-only, standard augmentation, and V3Cell augmentation), exactly as detailed in the Phenotyping augmentation classifier subsection. Each configuration was repeated with *n* = 5 independent random seeds sharing identical real-image splits and was evaluated on the fixed held-out real test set using macro-averaged *F*_1_. The mean and standard deviation across seeds were reported, and statistical comparison between augmentation conditions used paired-seed comparisons.

#### Experimental setup for fate prediction and trajectory modeling

This experiment evaluates both the per-sequence fate classifier and the full dynamic virtual 3D cell trajectory predictor on the longitudinal time-lapse dataset. The temporal model was trained on the 245 labeled sequences (28 cavity-formation, 106 cavity-collapse, 111 non-cavity) and evaluated on a separate, class-stratified held-out test set of *n* = 36 organoids (12 per fate class) that was disjoint from the 245 training sequences. The fate classifier was fine-tuned per the Stage 2 protocol above, and the optimal observation frame *k*^*\**^ was selected by 5-fold cross-validation on the 245 training sequences. Fate-prediction performance on the held-out test set was quantified by macro-averaged *F*_1_ and a per-class confusion matrix. The dynamic modeler (sequence encoder, MotionRNN and frozen amniotic-sac generator *G*_AS_) was jointly trained per the Stage 2 protocol and evaluated on the same held-out test set. Given the first *K* = 6 conditioning frames and the predicted fate *ĉ*, MotionRNN rolled out the remaining 19 frames to yield 25-frame predicted trajectories. Temporal fidelity was quantified by the per-frame fate-class probability on the ground-truth fate class, computed over the full 25-frame horizon, averaged across test organoids within each fate class, and smoothed by a Savitzky–Golay filter (window length 7, polynomial order 3). The same procedure was applied to the corresponding real frame sequences to obtain paired reference curves, and an unsupervised UMAP reducer fitted exclusively on real training-frame features was fixed and used to project held-out real and V3Cell-generated frames into the same embedding.

### 4.11 Quantification and statistical analysis

All quantitative analyses and statistical tests were performed in Python 3.10 using NumPy (1.26), SciPy (1.11), scikit-learn (1.4), scikit-image (0.22), umap-learn (0.5), and PyTorch (2.1). Deep-learning models were trained on NVIDIA A100 GPUs under CUDA 12.1. Microscopy image parsing and GLCM-based Haralick descriptor extraction were performed with scikit-image, and Savitzky–Golay smoothing was performed with SciPy. Random seeds were fixed for all stochastic procedures unless otherwise specified: the global seed was 42 for dataset splitting and UMAP projection, and *n* = 5 independent seeds {0, 1, 2, 3, 4} were used for the phenotyping augmentation study.

Sample sizes are reported in the relevant Methods subsection or figure legend. For distributional fidelity, KID and IS were computed over *n* = 10 independent sampling runs and reported as mean ± standard deviation. For phenotyping augmentation, macro-averaged *F*_1_ scores are reported as mean ± standard deviation across *n* = 5 seeds sharing identical physical training splits. For the fate prediction classifier, test-set performance was computed on *n* = 36 held-out organoids (12 per fate class), and per-frame fate-class probability trajectories were averaged over the same test set.

Two-sided Wilcoxon rank-sum tests [80] were used to compare distributions of Haralick descriptors and lineage-specific morphometrics between real and V3Cell-constructed organoids. Cohen’s *d* [66] quantified effect sizes, with |*d*| *<* 0.2 considered biologically negligible, 0.2 ≤ |*d*| *<* 0.5 small, 0.5 ≤ |*d*| *<* 0.8 moderate, and *d* ≥ |0.8| large. Statistical significance was set at *p <* 0.05 unless otherwise stated. Where multiple comparisons were performed within a single analysis, *p*-values were reported without additional multiple-testing correction because effect size (Cohen’s *d*) rather than *p*-value was used as the primary criterion for biological interpretation, following best-practice recommendations for large-sample comparisons where nominal significance is inflated by sample size. Unless otherwise specified, summary statistics in figures are reported as mean ± standard deviation. Temporal trajectory plots in Fig. 5C and Fig. 5G are presented as mean ± s.e.m. Box plots indicate median, interquartile range (IQR), and 1.5×IQR whiskers, with individual outliers plotted.

## Supplementary information

*Supplementary Note 1*. Stage 2 loss terms.

*Table S1*. Kernel Inception Distance and Inception Score for constructed virtual organoids from colon, lung, and stomach, reported as mean ± s.d. over 10 independent runs.

*Table S2*. Per-frame fate-classification performance on the held-out test set.

*Table S3*. Antibodies, isotype controls, and DAPI used for validation assays.

*Figure S1*. Complete micro-textural fidelity evaluation of virtual 3D cells across all organoid lineages and subtypes.

*Figure S2*. Extended lineage-specific morphometric evaluation.

*Figure S3*. Full-frame fate-conditioned virtual 3D cell trajectories.

*Figure S4*. Chemical-structure features do not separate toxic from non-toxic OPFRs.

*Figure S5*. Structure-based models cannot predict OPFR developmental toxicity at screening-relevant accuracy.

## Acknowledgements

The authors thank all members of the lab for their support.

## Funding

This work was supported in part by the National Key R&D Program of China under Grant 2025YFC3409000.

## Author contributions

Conceptualization, L.Y. and P.H. Methodology, L.Y., X.D., and P.H. Data collection, C.X. and J.H. Data curation, Z.Z. and P.C. Investigation, X.D. and P.H. Formal analysis, L.Y., X.D., C.X., J.H., and P.H. Writing—original draft, L.Y. and P.H. Writing—review & editing, L.Y., X.D., C.X., X.Z., Z.Z., P.C., X.Y., Z.Y., J.R., H.H., J.H., and P.H. Supervision, J.H. and P.H., and P.H. Funding acquisition, P.H.

## Competing interests

The authors declare no competing interests.

## Data availability

The data supporting the findings of this study are available at https://github.com/Laineyoulu/V3Cell.

## Code availability

The source code associated with this study is available at https://github.com/Laineyoulu/V3Cell.

## Notes

### Competing Interest Statement

The authors have declared no competing interest.

