## Supplementary Information for "V3Cell: A Vision-Guided Virtual 3D Cell Framework for Phenotypic Modeling and Perturbation Prediction"

### A Supplementary Figures

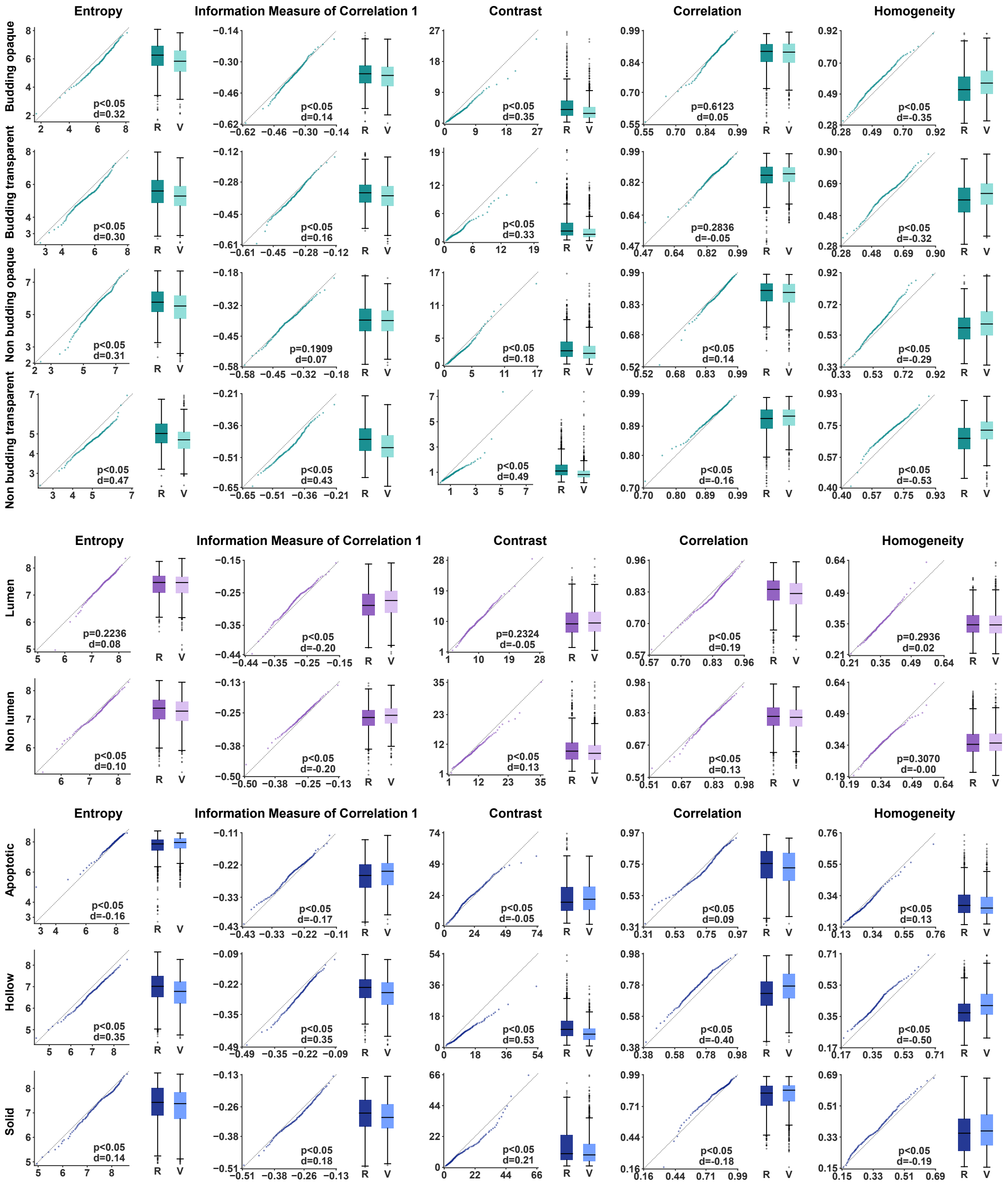

**Supplementary Fig. S1. Complete micro-textural fidelity evaluation of virtual 3D cells across all organoid lineages and subtypes.** Haralick texture descriptors (entropy, Information Measure of Correlation 1, contrast, correlation, and homogeneity) comparing real (R) and V3Cell-constructed virtual (V) samples. **(A)** Colon organoids: budding opaque, budding transparent, non-budding opaque, and non-budding transparent. **(B)** Lung organoids: lumen and non-lumen. **(C)** Stomach organoids: apoptotic, hollow, and solid. For each subtype–metric pair, quantile-quantile plots compare real and virtual distributions against the identity line. Paired box plots show marginal distributions. Wilcoxon rank-sum $p$-values and Cohen's $d$ effect sizes are annotated in each panel.

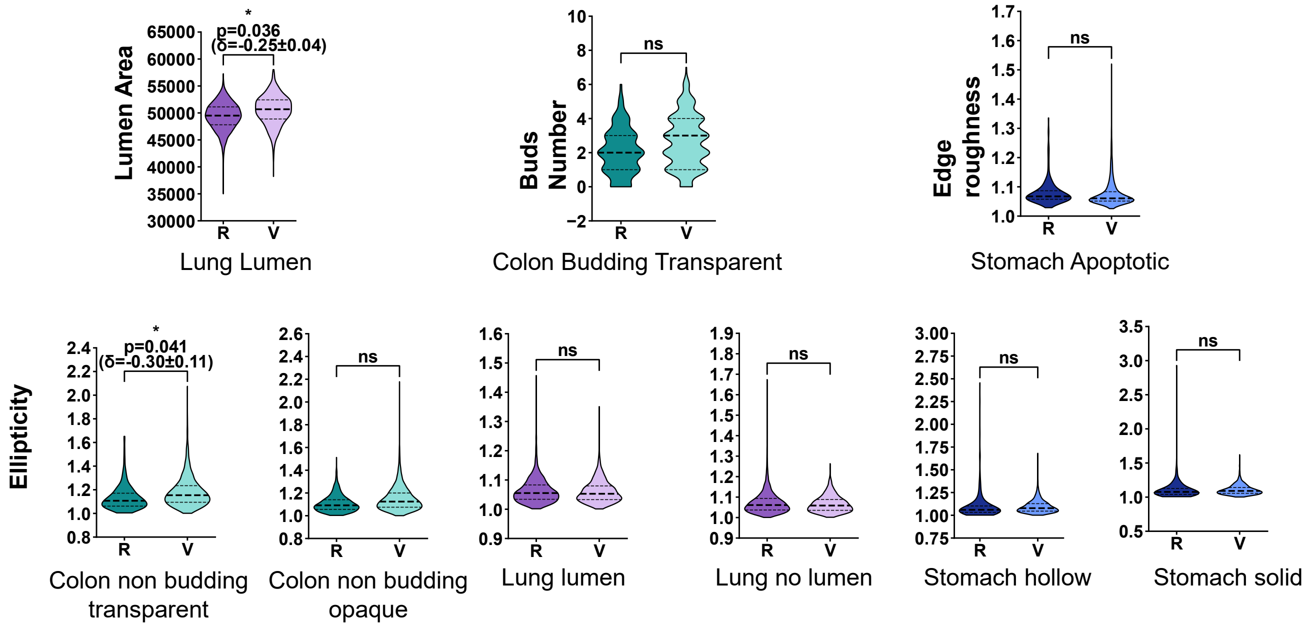

**Supplementary Fig. S2. Extended lineage-specific morphometric evaluation.** Comprehensive comparison of lineage-specific developmental hallmarks between real and V3Cell-constructed virtual organoids beyond the representative subtypes shown in the main text (Fig. 3E–F). Distributions of bud number (colon), lumen area (lung), and ellipticity (stomach) are shown for all evaluated subtypes within each lineage. Violin plots display paired real versus virtual distributions. Statistical significance was assessed by Wilcoxon rank-sum test, with effect sizes reported as Cohen's $d\pm$ s.e.

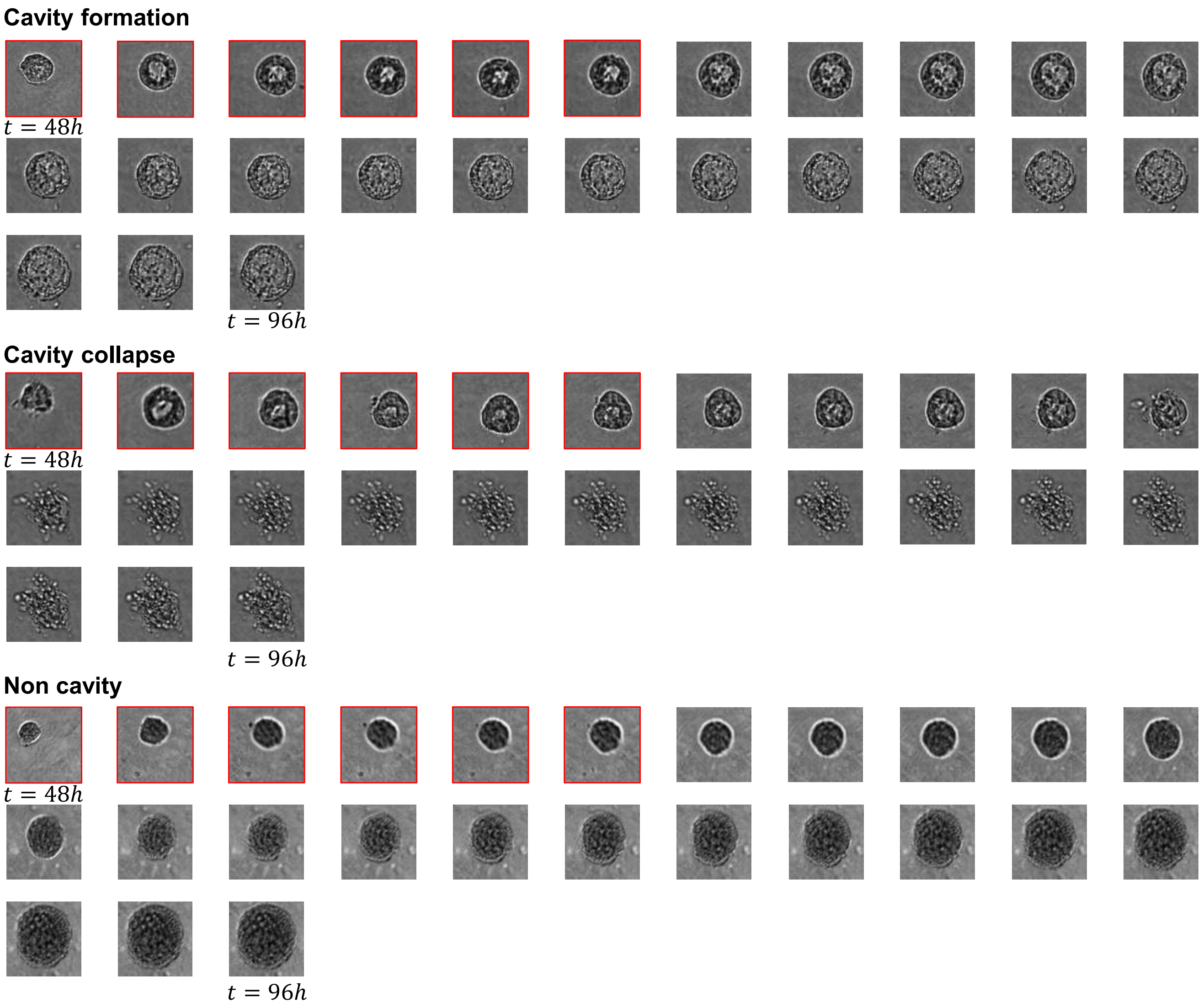

**Supplementary Fig. S3. Full-frame fate-conditioned virtual 3D cell trajectories.** Complete 25-frame predicted sequences for representative organoids from each of the three developmental fate classes: cavity formation (top), cavity collapse (middle), and non-cavity (bottom). Red-bordered frames indicate the $K = 6$ early-frame conditioning inputs at culture times $t = 48, 50, 52, 54, 56,$ and $58$ h. Subsequent frames are V3Cell-predicted outputs from $t = 60$ to $96$ h. Cavity-formation organoids progressively develop a central cavity surrounded by a rosette-like structure. Cavity-collapse organoids initially form a rosette but fail to maintain structural integrity, resulting in fragmentation. Non-cavity organoids grow as compact aggregates without cavity formation throughout the imaging window.

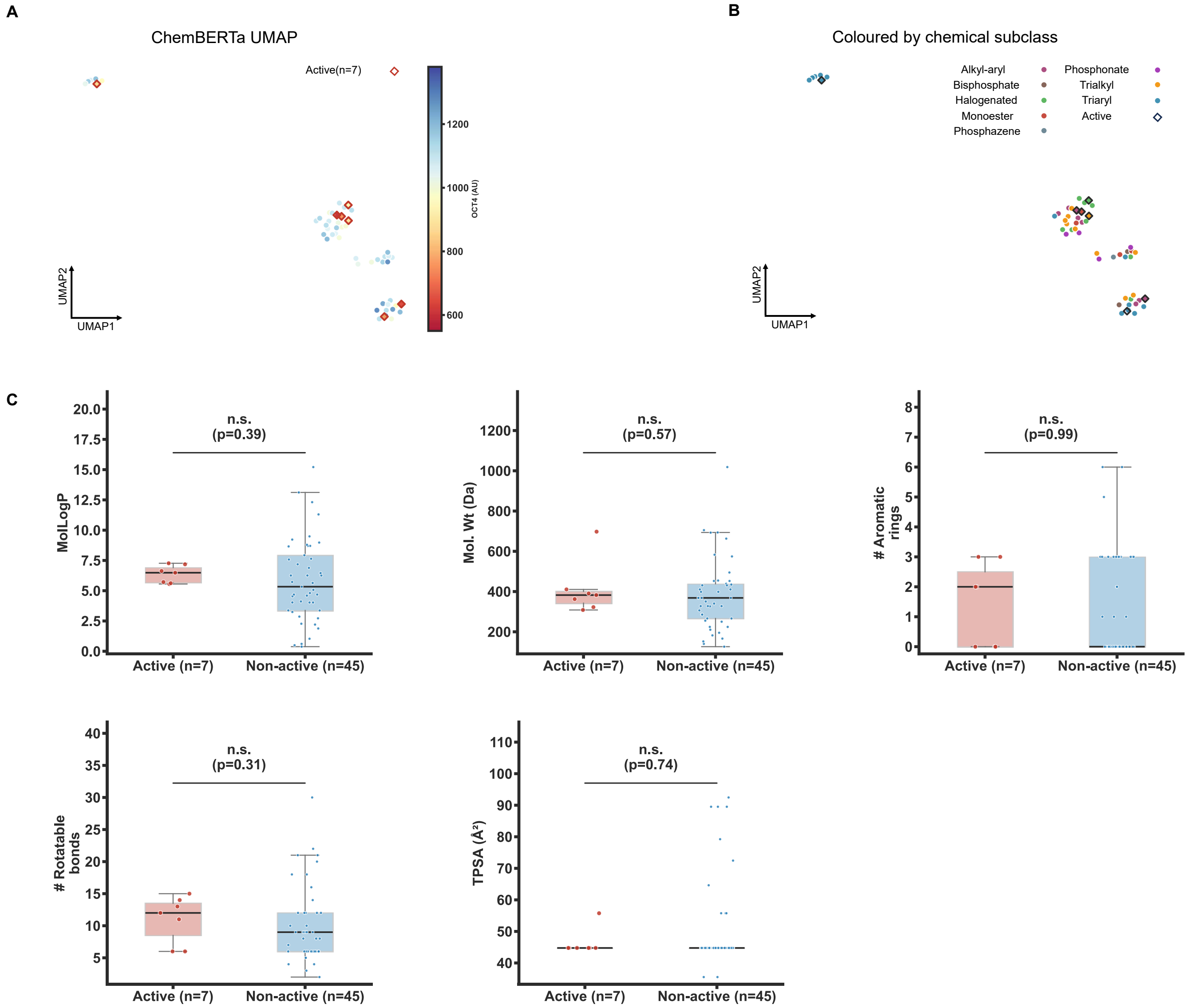

**Supplementary Fig. S4. Chemical-structure features do not separate toxic from non-toxic organophosphate flame retardants (OPFRs).** Analysis of 52 OPFRs (1 of 53 excluded, SMILES unavailable) against the OCT4 immunofluorescence developmental-toxicity readout from the source OPFR screen. The active label denotes OCT4 more than two standard deviations below the DMSO control (OCT4 $<$ 949 arbitrary units, $n = 7$ active, $45$ non-active). **(A)** Uniform Manifold Approximation and Projection (UMAP) of ChemBERTa molecular embeddings, colored by OCT4 intensity. Active compounds (diamonds) are dispersed across the embedding rather than forming a distinct cluster. **(B)** The same embedding colored by chemical subclass. Structure organizes by subclass, yet active compounds span multiple subclasses. UMAP projections are shown for illustration only. The quantitative assessment is in Supplementary Fig. S5. **(C)** Five physicochemical descriptors (calculated logP, molecular weight, number of aromatic rings, number of rotatable bonds, and topological polar surface area [TPSA]) for active versus non-active compounds. Boxes denote median and interquartile range. Points are individual compounds. No descriptor differs between groups (Mann–Whitney $U$ test, all $p > 0.30$, no multiple-comparison correction applied).

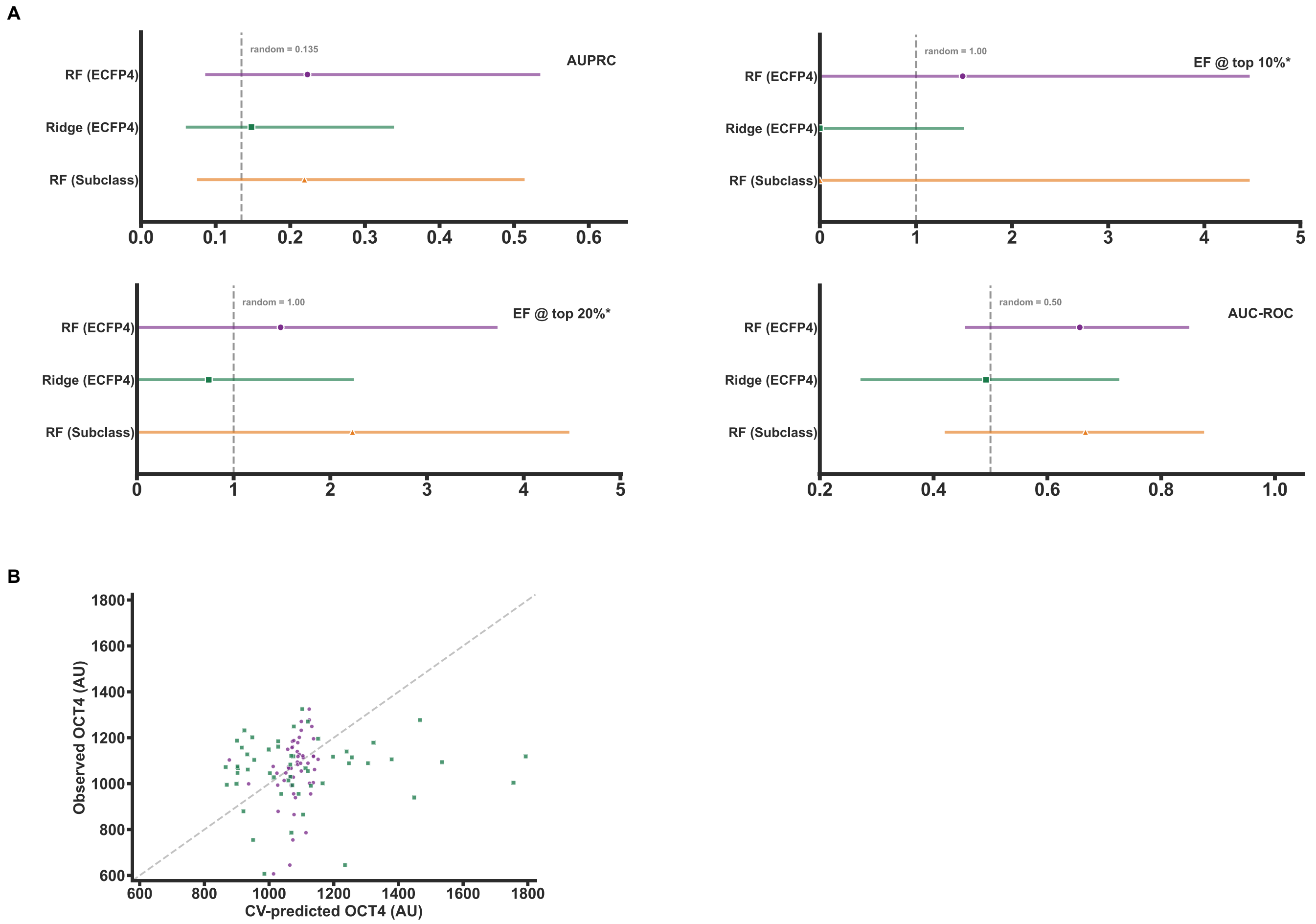

**Supplementary Fig. S5. Structure-based models cannot predict OPFR developmental toxicity at screening-relevant accuracy.** Prediction of the OCT4 readout from molecular structure for the 52 OPFRs in Supplementary Fig. S4, under scaffold-split (Bemis–Murcko) cross-validation following standard QSAR best practice. **(A)** Screening-grade metrics with 95% bootstrap confidence intervals ($n = 1000$) for a Random Forest (RF) and Ridge regression on ECFP4 fingerprints plus physicochemical descriptors, and for an RF trained on chemical-subclass labels alone: area under the precision–recall curve (AUPRC), enrichment factor (EF) among the top 10% and 20% of ranked compounds, and area under the receiver-operating-characteristic curve (AUROC). Dashed lines mark the random baselines (AUPRC $= 0.135$, the active-class prevalence. EF $= 1.0$. AUROC $= 0.50$). Every confidence interval includes the random baseline, and the subclass-only model matches the fingerprint model, indicating that the weak signal reflects coarse chemical-subclass enrichment rather than compound-level structure–activity relationships. Chemical subclasses were assigned from molecular structure, independent of the toxicity readout. **(B)** Observed versus cross-validated predicted OCT4 for RF (purple) and Ridge (green). Predictions do not track the identity line (dashed). Given the small number of actives ($n = 7$), we do not claim the absence of any structure–activity relationship in principle. Rather, no screening-relevant predictor can be built from this dataset, motivating the phenotype-based modeling used by V3Cell.

### B Supplementary Tables

**Supplementary Table S1.** Kernel Inception Distance (KID) and Inception Score (IS) for constructed virtual organoids across the colon, lung, and stomach lineages.

| Organ | Model | Runs | KID (↓) | IS (↑) |
|---|---|---|---|---|
| **Colon** | DiT | 10 | 0.000 066 ± 0.000 011 | 2.2258 ± 0.0492 |
| | Medfusion | 10 | 0.000 046 ± 0.000 001 | 2.2868 ± 0.0182 |
| | F2DGAN | 10 | 0.000 163 ± 0.000 001 | 2.4621 ± 0.0284 |
| | cGAN | 10 | 0.000 732 ± 0.000 001 | 1.1400 ± 0.0003 |
| | cVAE | 10 | 0.000 421 ± 0.000 001 | 1.7642 ± 0.0082 |
| | LDM | 10 | 0.001 074 ± 0.000 004 | 1.0094 ± 0.0012 |
| | V3Cell (Ours) | 10 | **0.000 019 ± 0.000 001** | **2.7734 ± 0.0229** |
| **Lung** | DiT | 10 | 0.000 046 ± 0.000 001 | 2.2983 ± 0.0218 |
| | Medfusion | 10 | 0.000 073 ± 0.000 002 | 2.2909 ± 0.0091 |
| | F2DGAN | 10 | 0.000 258 ± 0.000 002 | 2.2472 ± 0.0091 |
| | cGAN | 10 | 0.001 027 ± 0.000 002 | 1.0077 ± 0.0001 |
| | cVAE | 10 | 0.000 352 ± 0.000 002 | 1.7846 ± 0.0138 |
| | LDM | 10 | 0.000 959 ± 0.000 004 | 1.0266 ± 0.0019 |
| | V3Cell (Ours) | 10 | **0.000 017 ± 0.000 001** | **2.3122 ± 0.0224** |
| **Stomach** | DiT | 10 | 0.000 262 ± 0.000 002 | 2.2508 ± 0.0148 |
| | Medfusion | 10 | 0.000 100 ± 0.000 001 | 2.5030 ± 0.0110 |
| | F2DGAN | 10 | 0.000 082 ± 0.000 002 | 2.6305 ± 0.0237 |
| | cGAN | 10 | 0.001 019 ± 0.000 002 | 1.0625 ± 0.0004 |
| | cVAE | 10 | 0.000 515 ± 0.000 003 | 1.8573 ± 0.0060 |
| | LDM | 10 | 0.001 090 ± 0.000 007 | 1.0085 ± 0.0007 |
| | V3Cell (Ours) | 10 | **0.000 069 ± 0.000 002** | **2.8216 ± 0.0166** |

Note: Values are reported as mean ± s.d. over 10 independent runs. Bold indicates the best value within each organ system. Lower KID and higher IS indicate better distributional fidelity.

**Supplementary Table S2.** Per-frame fate-classification performance on the held-out test set.

| Frame $k$ | Accuracy | Macro-$F_1$ | Cavity-formation | Cavity-collapse | Non-cavity |
|---|---|---|---|---|---|
| 0 | 69.4% | 0.681 | 0.609 | 0.545 | 0.889 |
| 1 | 83.3% | 0.831 | 0.800 | 0.769 | 0.923 |
| 2 | 83.3% | 0.828 | 0.833 | 0.727 | 0.923 |
| 3 | 80.6% | 0.800 | 0.700 | 0.741 | 0.960 |
| **4** | **88.9%** | **0.888** | **0.870** | **0.833** | **0.960** |
| 5 | 80.6% | 0.800 | 0.737 | 0.741 | 0.923 |

Note: $n = 36$ held-out sequences (12 per fate class). Frame index $k$ runs over the six early conditioning frames, and $k^* = 4$ (bold) was selected by 5-fold cross-validation and used for fate prediction. Per-class values are $F_1$ scores.

**Supplementary Table S3.** Antibodies, isotype controls, and DAPI used for validation assays.

| Reagent | Company | Product number | Application and dilution |
|---|---|---|---|
| anti-OCT4 | CST | 75463 | 1:100 (IF) |
| anti-TFAP2A | Santa Cruz | sc12726 | 1:100 (IF) |
| anti-CDX2 | CST | 12306 | 1:100 (IF) |
| anti-Integrin $\beta_1$ | Abcam | ab24693 | 1:100 (IF) |
| anti-phospho-Myosin light chain (Ser19) | CST | 3671S | 1:50 (IF) |
| anti-rabbit IgG H&L | Abcam | ab150077 | 1:200 (IF) |
| anti-mouse IgG H&L | CST | 8890S | 1:200 (IF) |
| DAPI | Sigma | D9542 | 1:5000 (IF) |
| Alexa Fluor 488 Phalloidin | Molecular Probes | A12379 | 1:200 (IF) |
| anti-FAK | Abcam | ab40794 | 1:1000 (WB) |
| anti-phospho-FAK (Y397) | Abcam | ab81298 | 1:1000 (WB) |
| anti-GAPDH | CST | 3700S | 1:2000 (WB) |
| anti-Mouse IgG HRP-linked | CST | 7076S | 1:5000 (WB) |
| anti-Rabbit IgG HRP-linked | CST | 14708S | 1:1000 (WB) |
| Rabbit IgG | Abcam | ab125938 | – |
| Mouse IgG1 | Abcam | ab280974 | – |
| Mouse IgG2b | Abcam | ab281590 | – |

Note: IF, immunofluorescence. WB, Western blot. Dashes denote isotype controls used without a fixed working dilution.

### C Supplementary Note 1: Stage 2 loss terms

The composite Stage 2 objective is defined in the main text (Methods, Dynamic virtual 3D cell modeler). The reconstruction term $\mathcal{L}_{\text{recon}}$ and the adversarial terms $\mathcal{L}_{\text{adv}}^{\text{2D}}$ and $\mathcal{L}_{\text{adv}}^{\text{3D}}$ are given there. The remaining terms are defined below. Here, $B$ is the batch size, $T$ is the sequence length, $K$ is the number of conditioning frames, and $n = T - K$ is the number of autoregressively generated steps.

**Feature matching ($\mathcal{L}_{\text{feat}}$).**

$$\mathcal{L}_{\text{feat}} = \frac{1}{N_D} \sum_{i=1}^{N_D} \sum_{j=1}^{L_i - 1} \left\| \operatorname{sg}\big(\phi_j^i(\mathbf{x})\big) - \phi_j^i(\hat{\mathbf{x}}) \right\|_1, \tag{S1}$$

where $N_D = 2$ is the number of discriminator scales, $\phi_j^i$ is the activation at layer $j$ of scale $i$, and the final scalar-output layer is excluded. The operator $\operatorname{sg}(\cdot)$ stops gradients through the real-video features.

**Contrastive loss ($\mathcal{L}_{\text{contrastive}}$).** The first real frame is the anchor and a randomly sampled generated frame from the same sequence is the positive sample. Features are encoded by the ContrastiveEncoder and $L_2$-normalized. After the first-in–first-out memory bank of $K_{\text{mb}} = 4{,}096$ historical real-frame features is populated,

$$\mathcal{L}_{\text{contrastive}} = -\frac{1}{B} \sum_{i=1}^{B} \log \frac{\exp(s_i^{\text{pos}}/\tau)}{\exp(s_i^{\text{pos}}/\tau) + \sum_{k=1}^{K_{\text{mb}}} \exp(s_{i,k}^{\text{neg}}/\tau)}, \qquad \tau = 0.07, \tag{S2}$$

where $s$ denotes cosine similarity. Before the memory bank is populated, the other in-batch samples serve as negatives.

**Temporal direction loss ($\mathcal{L}_{\text{dir}}$).** With per-frame pixel differences $\Delta\hat{\mathbf{x}}_t = \hat{\mathbf{x}}_{t+1} - \hat{\mathbf{x}}_t$ and $\Delta\mathbf{x}_t = \mathbf{x}_{t+1} - \mathbf{x}_t$, flattened over $C \times H \times W$,

$$\mathcal{L}_{\text{dir}} = \frac{1}{B(T-1)} \sum_{b=1}^{B} \sum_{t=1}^{T-1} \left( 1 - \frac{\Delta\hat{\mathbf{x}}_t^{(b)} \cdot \Delta\mathbf{x}_t^{(b)}}{\|\Delta\hat{\mathbf{x}}_t^{(b)}\|_2 \, \|\Delta\mathbf{x}_t^{(b)}\|_2} \right). \tag{S3}$$

This term is evaluated over the full sequence, including conditioning and predicted frames.

**Mutual-information loss ($\mathcal{L}_{\text{MI}}$).** For each generated step $t = K+1, \ldots, T$, the injected noise $\boldsymbol{\epsilon}_t \sim \mathcal{N}(\mathbf{0}, \mathbf{I})$ is reconstructed as $\hat{\boldsymbol{\epsilon}}_t$ from the MotionRNN hidden state. Flattening over batch and the $n$ generated steps, with $N = Bn$,

$$\mathcal{L}_{\text{MI}} = -\frac{1}{N} \sum_{i=1}^{N} \frac{\hat{\boldsymbol{\epsilon}}_i \cdot \operatorname{sg}(\boldsymbol{\epsilon}_i)}{\|\hat{\boldsymbol{\epsilon}}_i\|_2 \, \|\operatorname{sg}(\boldsymbol{\epsilon}_i)\|_2}, \tag{S4}$$

where $\operatorname{sg}(\cdot)$ stops gradients through the sampled noise.

**Motion smoothness loss ($\mathcal{L}_{\text{motion}}$).** With motion-latent increments $\Delta \mathbf{z}_t = \mathbf{z}_t^M - \mathbf{z}_{t-1}^M$ over the $n$ generated steps and motion-latent dimension $d_M$,

$$\mathcal{L}_{\text{motion}} = \frac{1}{B\, n\, d_M} \sum_{b,t,j} \big(\Delta z_t^{(b,j)}\big)^2 + \frac{1}{2}\, \frac{1}{B(n-1)d_M} \sum_{b,t,j} \big(\Delta z_{t+1}^{(b,j)} - \Delta z_t^{(b,j)}\big)^2. \tag{S5}$$

The first term penalizes increment magnitude and the second term penalizes second-order temporal changes.

**KL regularization ($\mathcal{L}_{\text{KL}}$).** The encoder posterior $q_\phi(\mathbf{z} \mid \mathbf{x}) = \mathcal{N}(\boldsymbol{\mu}, \boldsymbol{\sigma}^2)$ is regularized toward the standard Gaussian prior:

$$\mathcal{L}_{\text{KL}} = \frac{1}{2} \sum_j \big(\mu_j^2 + \sigma_j^2 - \log \sigma_j^2 - 1\big). \tag{S6}$$
